# Structure and mechanistic analyses of the gating mechanism of elongating ketosynthases

**DOI:** 10.1101/2021.02.13.431092

**Authors:** Jeffrey T. Mindrebo, Aochiu Chen, Woojoo E. Kim, Rebecca N. Re, Tony D. Davis, Joseph P. Noel, Michael D. Burkart

**Author notes:** These authors contributed equally to this work (Jeffrey T. Mindrebo, Aochiu Chen). **Author Contributions**: J.P.N. and M.D.B. supervised the project. J.T.M and A.C. designed the project. J.T.M. and A.C. performed crystallographic studies. J.T.M. carried out kinetic assays. A.C. and W.E.K performed crosslinking studies. W.E.K, R.N.R, and T.D.D. synthesized and characterized probe molecules. J.T.M., A.C., J.P.N, and M.D.B. wrote and edited the manuscript.

## Abstract

Ketosynthases (KSs) catalyze carbon-carbon bond forming reactions in fatty acid synthases (FASs) and polyketide synthases (PKSs). KSs utilize a two-step ping pong kinetic mechanism to carry out an overall decarboxylative thio-Claisen condensation that can be separated into the transacylation and condensation reactions. In both steps, an acyl carrier protein (ACP) delivers thioester tethered substrates to the active sites of KSs. Therefore, protein-protein interactions (PPIs) and KS-mediated substrate recognition events are required for catalysis. Recently, crystal structures of *Escherichia coli* elongating type II FAS KSs, FabF and FabB, in complex with *E. coli* ACP, AcpP, revealed distinct conformational states of two active site KS loops. These loops were proposed to operate via a gating mechanism to coordinate substrate recognition and delivery followed by catalysis. Here we interrogate this proposed gating mechanism by solving two additional high-resolution structures of substrate engaged AcpP-FabF complexes, one of which provides the missing AcpP-FabF gate-closed conformation. Clearly defined interactions of one of these active site loops with AcpP are present in both the open and closed conformations, suggesting AcpP binding triggers or stabilizes gating transitions, further implicating PPIs in carrier protein-dependent catalysis. We functionally demonstrate the importance of gating in the overall KS condensation reaction and provide experimental evidence for its role in the transacylation reaction. Furthermore, we evaluate the catalytic importance of these loops using alanine scanning mutagenesis and also investigate chimeric FabF constructs carrying elements found in type I PKS KS domains. These findings broaden our understanding of the KS mechanism which advances future engineering efforts in both FASs and evolutionarily related PKSs.

## Introduction

Fatty acid biosynthesis (FAB) is an essential primary metabolic pathway conducive to metabolic engineering.^1–10^ Fatty acid synthases (FAS) carry out biosynthesis by iteratively condensing and reducing malonyl-CoA derived 2-carbon units.^1–2^ FAS can be organized as either large, single gene, multidomain megasynthases, type I, or as discrete, multiple gene, monofunctional enzymes, type II.1.3,11-13 Despite differences in structural organization, the biochemistry of type I and II FAS are generally conserved.^2^ Additionally, type II FASs are considered to be the evolutionary progenitors of polyketide synthases (PKSs), a class of enzymes known to produce a variety of structurally complex and biologically active compounds.^14–18^ Ketosynthases (KSs) initiate each round of FAS by condensing a growing acyl-chain with 2-carbon units via a decarboxylative Claisen condensation with the 3-carbon malonyl-CoA substrate to produce a β-keto intermediate. Subsequent reactions catalyzed by the ketoreductase (KR), dehydratase (DH), and enoylreductase (ER) fully reduce the β-carbon before another round of chain extension or offloading of the mature fatty acid (FA). Central to this process is the small 9 kDa acyl carrier protein (ACP). The ACP carries thioester-tethered pathway intermediates to each enzyme active site using a post-translationally installed 4’-phosphopantetheine arm (PPant).^19–20^ ACPs must form transient, productive protein-protein interactions (PPIs) with each respective partner enzyme (PE) in order to deliver their substrates in catalytic competent forms.^21–30^

KS-mediated carbon-carbon bond formation is the primary driving force of FAB. These enzymes use a two-step kinetic ping-pong bi-bi mechanism that can be viewed mechanistically as transacylation and condensation reactions (Figure 1A). ^30–33^ In the transacylation step, acyl-ACP binds to the KS and transfers its thioester-bound cargo to the KS active site cysteine residue, producing an acyl-KS thiointermediate.^34–36^ In the condensation step, malonyl-ACP associates with the KS and undergoes decarboxylation to produce a keto-enolate tautomerization, wherein the carbanion tautomer condenses with the thioester-bound FA to produce the β-ketoacyl-ACP product. Mechanistic elements of the transacylation and condensation steps are still contested, and a detailed molecular understanding for KS substrate specificity is therefore lacking.^31–33^ FA chain extension is energetically coupled to decarboxylation, making the forward reaction thermodynamically favored and essentially irreversible upon loss of CO2. Therefore, KSs play a central role in controlling metabolic flux and FA chain length.^4–31^ This point has been well demonstrated by recent successes in rationally retooling the product profile of the fungal FAS.^6–37–39^ Future engineering efforts, in both FAS and PKS platforms, will benefit from a more in depth understanding of the reaction mechanisms and substrate specificities of functionally diverse KSs.^40–41^

**Figure 1:**
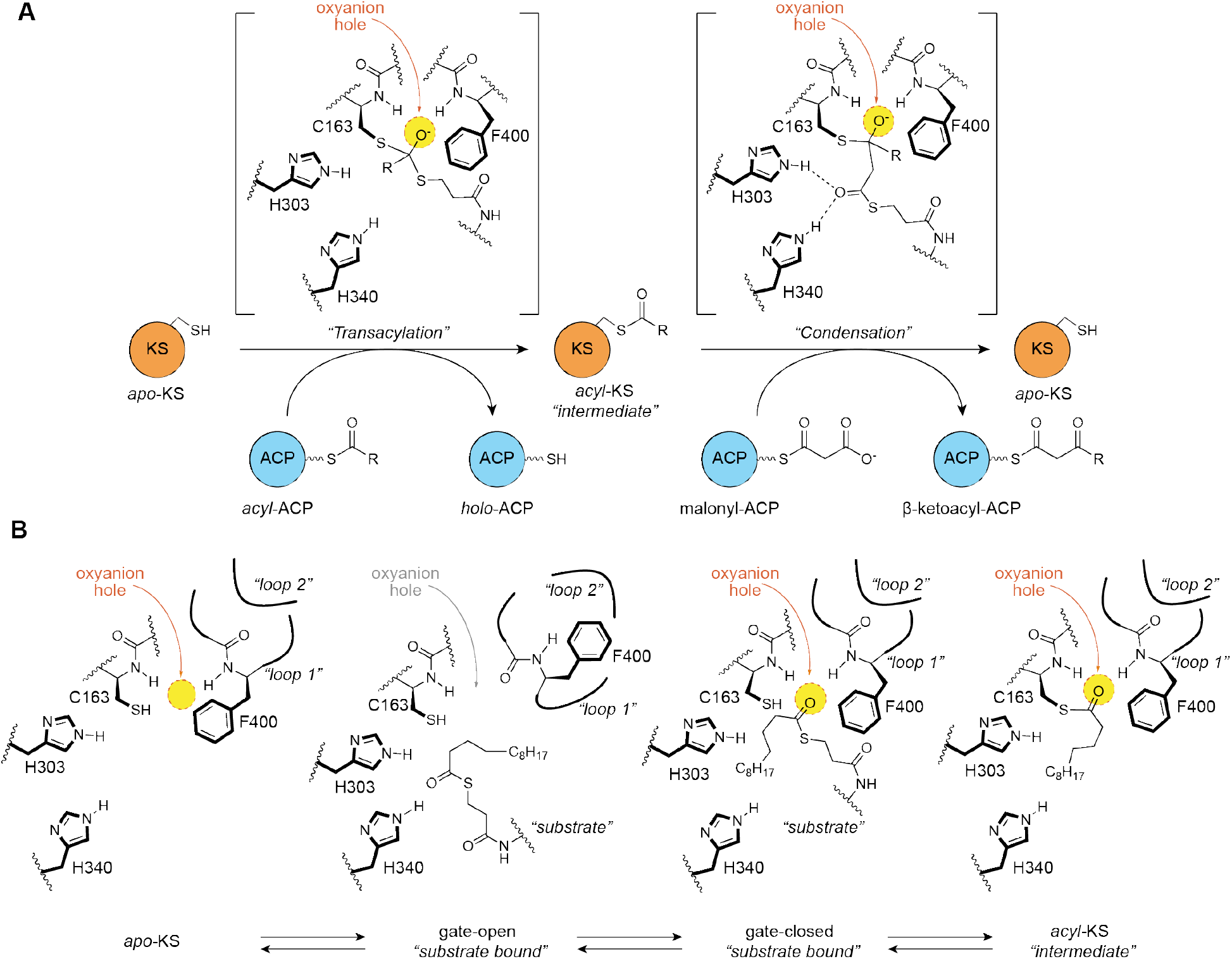
KS ping-pong reaction and gating mechanism overview. A) Ketosynthase ping-pong mechanism with active site schematic showing oxyanion hole (yellow circle) stabilized tetrahedral intermediates that are formed during the transacylation and condensation half-reactions. B) Proposed ketosynthase gating mechanism coordinated by loops 1 and 2 and the Phe400 gating residue. Each active site represents a step forward during the transacylation half-reaction starting with the *apo-*KS active site and ending with the acyl-KS intermediate. (TWO COLUMN)

*Escherichia coli* type II FAS has long served as a model system for understanding the structure, function, and mechanism of FASs.^1–42–43^ *E. coli* possesses two related elongating KSs, FabB and FabF, that have broad overlapping substrate specificities and are representative of specific KS families, KASI and KASII, respectively.^44–45^ The active sites of elongating KSs are comprised of a Cys-His-His catalytic triad, where the cysteine, positioned at the N-terminus of a long α-helix, is the active site nucleophile and the two histidines participate in the decarboxylation step. The oxyanion hole formed by the backbone amides of the catalytic cysteine (Cys163) and a highly conserved phenylalanine gating residue (Phe400) stabilize oxyanion tetrahedral intermediates formed during both half-reactions (Figure 1A).^34–36^ Additionally, Phe400 is known to regulate access to the active site by sensing the acylation state of the catalytic cysteine.^32–34–36^

In our previous work, we determined crystal structures of FabB and FabF crosslinked to *E. coli* ACP (AcpP) loaded with either C12- or C16-α-bromo-pantetheinamide substrate mimetics.^24^ These structures, along with biochemical and computational studies, led to the proposal of a double drawbridge-like gating mechanism comprised of two active site loops, loop 1 and loop 2, that undergo conformational rearrangements to regulate KS substrate recognition and processing (Figure 1B). During substrate delivery, the gate opens to allow access to the KS active site, and in doing so, disrupts the oxyanion hole (Figure 1B). The active site machinery only reforms upon gate closure, thereby providing a means to regulate selectivity by organizing the oxyanion hole for acyl transfer when the correct substrate is bound. Loop 1 consists of a highly conserved GFGG motif, which includes the conserved phenylalanine gating residue, Phe400. Loop 2 is distal to the active site and abuts loop 1 in the gate-closed conformation. A conserved loop 2 aspartate residue, Asp265, stabilizes the gate-open state by coordinating a complex hydrogen-bonding interaction network with loop 1 (Figure 2A). Interestingly, much of the loop 2 motif appears to be only conserved within, but not between, different KS families (Figure 2B).

**Figure 2:**
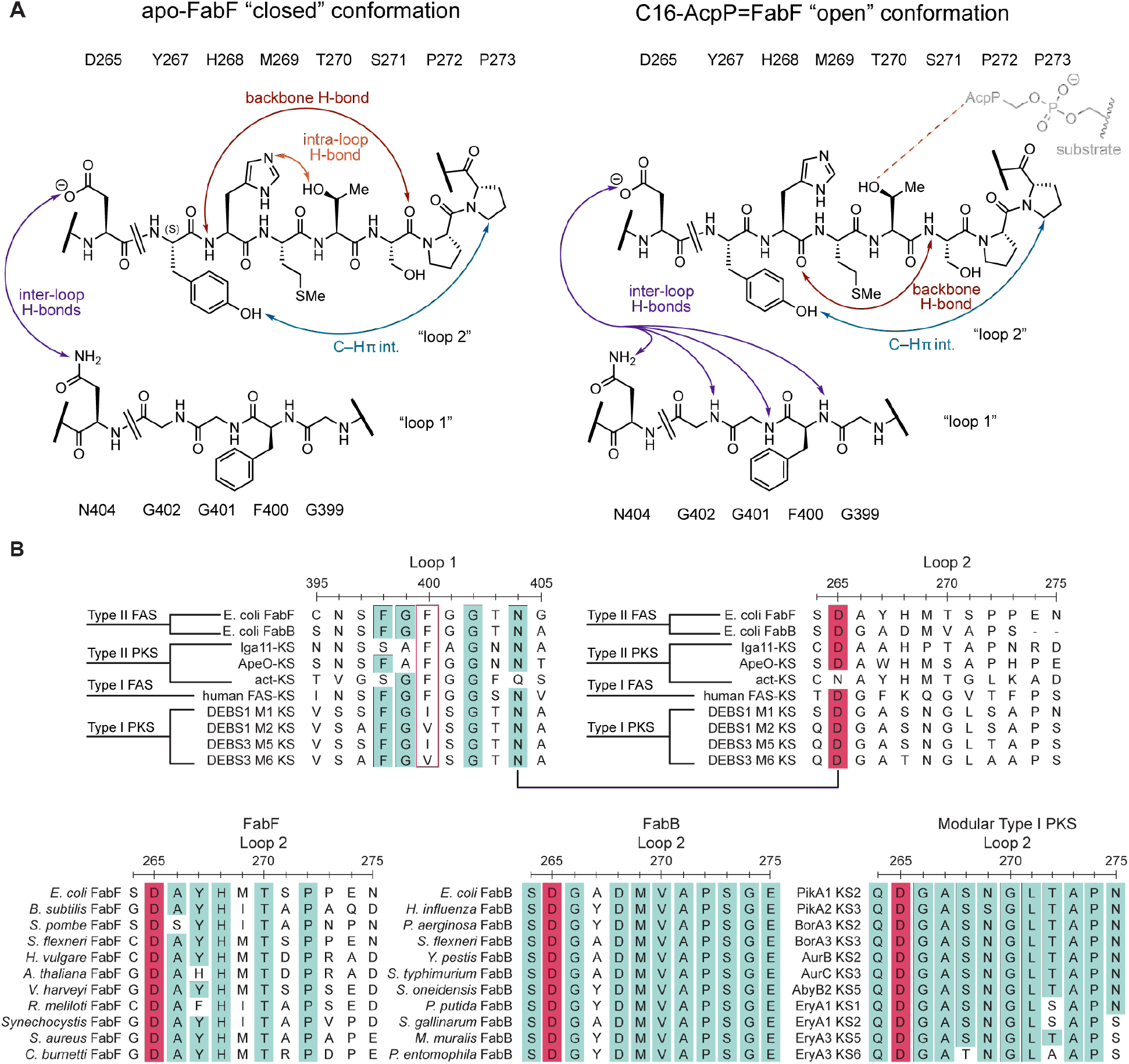
Gating loop interaction network and conservation. A) Schematic for loop 1–loop 2 interactions in the closed (*apo-*FabF) and open (C16-AcpP=FabF) states. Interactions between specific residues are shown by arrows. FabF loop 2 sequence is above the loop 2 peptide schematic and the FabF loop 1 sequence is below the schematic. B) Sequence alignment between FabF gating loops with the putative gating elements found in different classes of condensing enzymes. The loop 1 gating residue (Phe400) is outlined in a red box and the interaction between N404 and the conserved D265 is indicated by the connecting line segment. Loop 2 sequences from within 3 different classes of condensing enzymes (FabF, FabB, and modular type I PKS) are shown in separate alignments highlighting the conservation within, but not between, families of condensing enzymes. All residues above an identity threshold of 70% are highlighted.

Currently, only the AcpP-bound gate-open conformation is available for FabF, while only the AcpP-bound gate-closed conformation is available for FabB, making it difficult to directly compare the open and closed conformations in the same condensing enzyme family. Additionally, catalytic evidence that the proposed gating mechanism does indeed affect substrate turnover, and more specifically, the transacylation step, has not yet been demonstrated. In the work reported herein, we use mechanistic crosslinkers carrying different substrate mimetics to elucidate x-ray crystal structures of two additional acyl-AcpP=FabF (where the ‘=’ denotes a crosslink) complexes, one of which is in the missing gate-closed conformation. Next, we employ two HPLC-based kinetic assays to validate the importance of the KS gating machinery for the overall KS condensation reaction and transacylation half-reaction. We then explore the role of loop 2 using alanine scanning mutagenesis and demonstrate its conformational relevance to enzyme catalysis, providing additional evidence for an allosteric connection between AcpP binding and gating events. Finally, we characterize a panel of FabF-Type I PKS KS loop 1 chimeras, where FabF’s GFGG motif is replaced by corresponding GVSG and GISG motifs found in the *cis-AT* modular type I PKS KS domains. Results from our study provide the missing AcpP-FabF gate-closed conformation structure as well as structural and biochemical support for the importance of gating in type II FAS KSs. These findings have broad implications for KS-directed metabolic engineering in FAS and provide additional insights into how these putative gating elements may operate in the evolutionarily related PKS condensing enzyme families.

## Results and Discussion

### Development and analysis of a C16:1-α-bromo-pantetheinamide crosslinking probe

In order to compare the open and closed states within a single family of condensing enzymes, we aimed to crystallize an acyl-AcpP=FabF complex in a catalytically relevant, gate-closed conformation. We first developed a pantetheinamide mechanistic crosslinker that we reasoned could trap the AcpP-FabF complex in a closed conformation. Previous studies regarding FabF’s substrate specificity show that, unlike FabB, FabF is responsible for elongation of *cis*-palmitoeloyl-ACP (C16:1) to *cis*-vaccenoyl-ACP (C18:1). While *cis*-vaccenate is not essential for *E. coli* survival, this FabF product is responsible for *E. coli’s* rapid homeoviscous adaptive response to regulate membrane fluidity in response to reductions in temperature.^44–48^ Therefore, we developed a synthetic route for a C16:1-α-bromo-pantetheinamide unsaturated crosslinking probe (Figure 3), designed to mimic FabF’s privileged substrate, palmitoleoyl-ACP (Figure S1).

**Figure 3:**
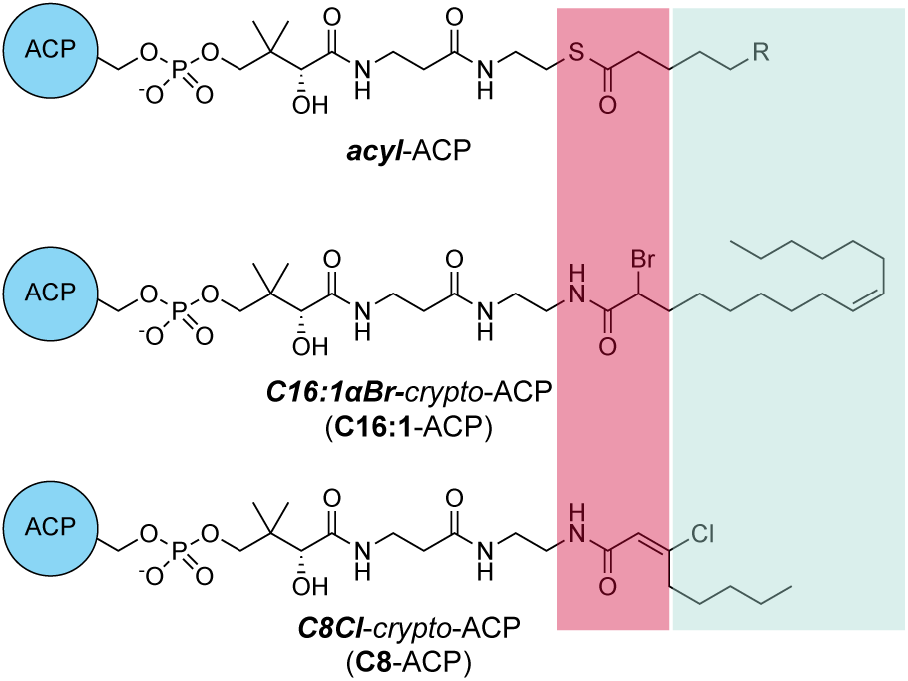
Crosslinking probes. Comparison of acyl-AcpP with crosslinker-loaded *crypto-AcpPs* utilized in this study to trap and crystalize AcpP=FabF complexes. An acyl-ACP substrate is shown to demonstrate the chemical structure of the natural substrate. The two crosslinking pantetheinamide probes have an amide instead of a thioester linking their acyl substrates as well as either an α-bromo or chloroacrylamide warhead. The region that has been modified to facilitate stable crosslinkers is shown in red while the substrate mimetics are outlined in light green.

Using a one-pot chemoenzymatic method,^49^ we loaded the C16:1-α-bromopantetheinamide crosslinking probe onto *apo-*AcpP to produce C16:1αBr-*crypto-*AcpP (C16:1-AcpP) (Figure 3, Figure S2). We then tested the crosslinking efficiency and rate of C16:1-AcpP with wt FabF using a time course gel-based crosslinking assay. Similar to previously tested α-bromo crosslinkers, C16:1-AcpP crosslinks with FabF to completion in under ten minutes across a range of pHs (Figure S3). Synthesis of different chain length, unsaturated crosslinkers may aid in elucidating structures of ACP-partner enzyme complexes with additional metabolic enzymes, such as *E. coli* FabB^50–52^ or the FatA and FatB FAS thioestersases^53–55^ from plant plastids, to elucidate their respective substrate specificities and recognition mechanisms.

### C16:1-AcpP=FabF captures distinct substrate conformation

Cross-seeding using C16-AcpP=FabF^24^ crystals nucleated diffraction-quality crystals of C16:1-AcpP=FabF for x-ray data collection, structure determination and refinement to a nominal resolution of 2.0-Å (Table S1, Figure S4). The complex crystallized in the same space group and unit cell as that of C16-AcpP=FabF (PDB ID: 6OKG), with an asymmetric unit containing a FabF monomer crosslinked to a single AcpP. Structural alignment with C16-AcpP=FabF (PDB ID: 6OKG) shows that these two complexes are nearly identical and overlay with an RMSD of 0.124 Å. The gating loops in the C16:1AcpP=FabF complex are in the open conformation, indicating that the gate-open conformation is favored in the presence of a crosslinked AcpP bearing long-chain α-bromo crosslinkers. (Figure 4A-C, Figure S5)

**Figure 4:**
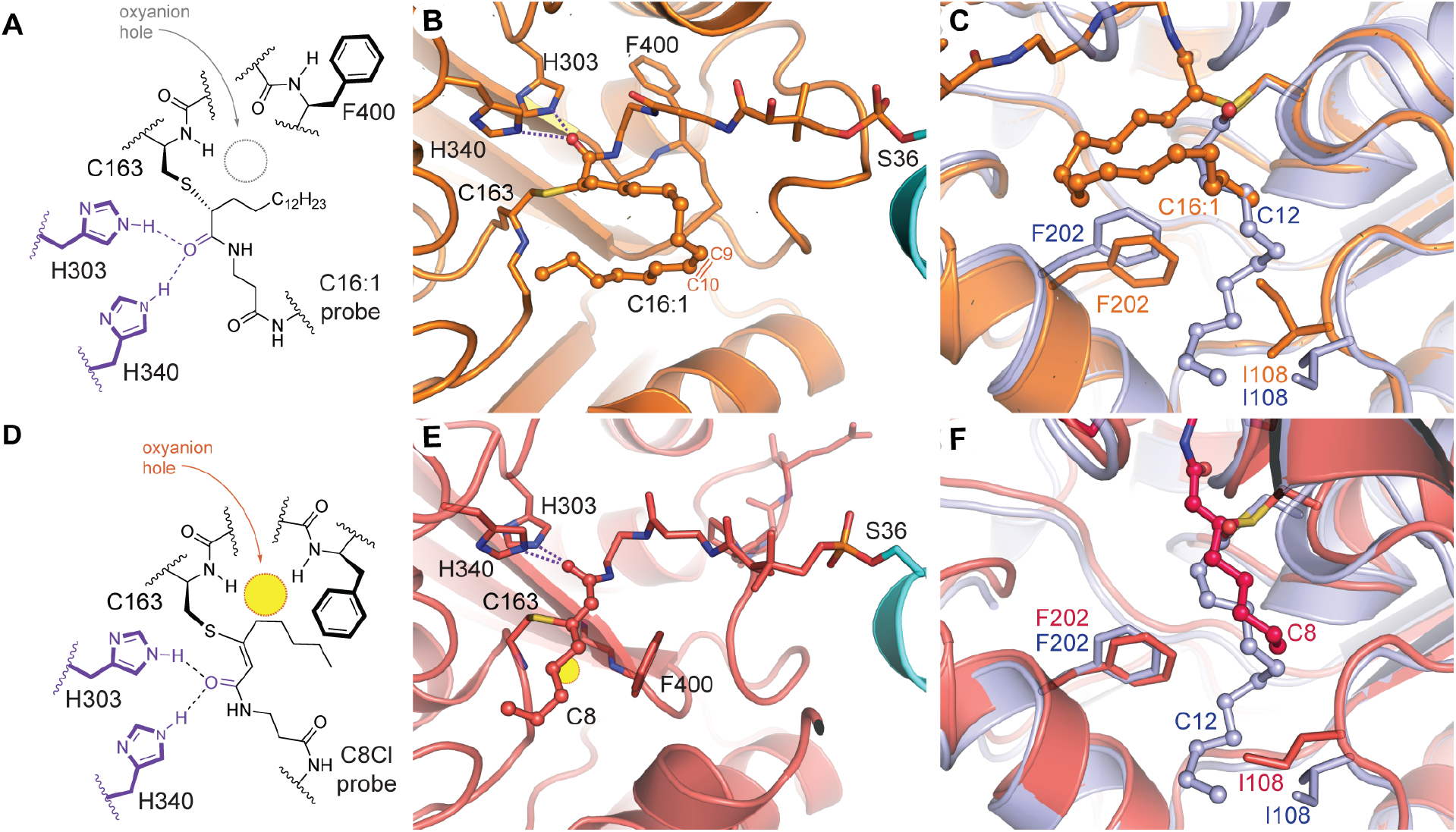
C16:1-and C8-AcpP=FabF active site organization. A) 2D schematic of C16:1-AcpP=FabF active site. The oxyanion hole is not formed and Phe400 is rotated away from the active site. B) 3D rendering of C16:1-AcpP=FabF active site and crosslinker. C) Acyl-binding pocket of C16:1-AcpP=FabF compared to that of C12-FabF (PDB: 2GFY). D) 2D schematic of C8-AcpP=FabF active site. The oxyanion hole is formed and Phe400 and gating loops are in the closed conformation. E) 3D rendering of C8-AcpP=FabF active site and crosslinker. F) Acyl-binding pocket of C8-AcpP=FabF compared to that of C12-FabF (PDB: 2GFY).

Closer analysis of the crosslinker indicates that the stereocenter of the α-carbon at the thioether crosslink is in the *R* configuration, while that of the previously solved structures were assigned to the *S* configuration (Figure 4A, Figure S6). In the C16-AcpP=FabF complex, the C16 acyl chain extends away from the active site, whereas in C16:1-AcpP=FabF, the *cis*-double bond of the unsaturated probe is in a kinked conformation that redirects the polymethylene chain toward the acyl-binding pocket. (Figure 4B-C, Figure S5) The terminal methyl moiety of the acyl chain is positioned in front of the back gate, Ile108 and Phe202, of the acyl-binding pocket (Figure 4C).^35–56^ When compared to the C12-bound acyl-enzyme intermediate structure of FabF (C12-FabF, PDB: 2GFY), the Ile108-Phe202 gate is in a closed conformation (Figure 4C). Despite the closed back gate, the tail of the kinked alkyl substrate is positioned to enter the acyl binding pocket. This may indicate that the Ile108-Phe202 back gate of FabF, which is replaced by glycine in FabB, plays a role in substrate-intermediate processing. Previous studies have shown that altering the identity of this gating residue results in antibiotic resistance^57–58^ and changes in KS chain length specificity in both bacterial and fungal FAS systems.^37–58^

The kinked C16:1 substrate favors a more compact conformation, which may be more readily accommodated and processed compared to the saturated C16 substrate. These observations, along with differences in the acyl-binding pocket and divergent loop 2 sequence found in the FabF condensing enzyme family, may explain FabF’s substrate preference for C16:1 over C16.^44–45^ Future mutagenic work probing the Ile108-Phe202 back gate and/or loop 2 may provide insights into FabF’s cryptic role in homeoviscous adaptation.

### C8-AcpP=FabF structure provides an ACP-bound gate-closed conformation

AcpP loaded with the *trans*-C8-chloroacrylate PPant probe, C8Cl-*crypto-*AcpP (C8-AcpP) (Figure 3), requires several hours to crosslink to FabF, but when loaded with an α-bromo crosslinker, such as the C16:1-α-bromo probe in this study, AcpP crosslinks in minutes.^24^ This significant difference in crosslinking rates and the propensity for α-bromo crosslinkers to trap FabF in the gate-open conformation led us to hypothesize that crosslinking to *trans*-C8-chloroacrylate may require a slow, stochastic conformational change to the closed form to react with the active site cysteine of FabF. Therefore, we obtained and collected data on diffraction quality crystals of C8Cl-*crypto-*AcpP=FabF (C8-AcpP=FabF), which upon structure elucidation and refinement, resulted in a 2.65-Å-resolution structure. The asymmetric unit of C8-AcpP=FabF contains the FabF dimer crosslinked to two AcpPs. Analysis of the active sites shows reliable electron density for the PPant arm and the thiovinyl covalent crosslink between C3 and Cys163 (Figure S7). The gating loops are positioned in the catalytically competent gate-closed conformation and an organized, but unoccupied, oxyanion hole is formed by the backbone amides of Cys163 and Phe400 (Figure 4D,E). The PPant moiety of the probe is anchored by polar contacts with Thr270, Ser271, Thr305, and Thr307. The carbonyl oxygen of the substrate mimetic coordinates to His303 and His340, with the latter histidine serving an essential role for the condensation reaction.^32–33^ Similar probe-KS interactions were observed in the crystal structures of FabB-AcpP^23–24^ and IgaKS-IgaACP^59^, indicating an evolutionarily conserved PPant binding site.

The acylated C12-FabF structure (PDB: 2GFY) overlays with the KS domain from C8Cl-AcpP=FabF with an RMSD of 0.314 Å (Figure 4F). The active sites of these complexes are in nearly identical conformations, including the gating loops, with the Phe400 gating residue rotated and translated away from Cys163 to form the presumptive malonyl-AcpP binding pocket.^34–35–60^ Interestingly, the chlorovinyl probe has the same carbon count as the putative condensation intermediate. The carbonyl group of the FA substrate mimetic interacts with His303 and His340, positioning the C2 carbon in close proximity to the Cys163-bound thioester carbonyl carbon (Figure 5). With the gating loops closed, the oxyanion hole is organized, which, as seen when overlaid with C12-FabF, would be occupied by the carbonyl oxygen of the bound FA. Even with the altered Cys163 position in C8-AcpP=FabF compared to C12-FabF, angle measurements in the overlay place the C2 carbon within 30 degrees of the Burgi-Dunitz angle (ca. 105°) (Figure 5). Given that the observed probe behavior matches the current knowledge of the condensation mechanism, we propose that the C8 chlorovinyl probe mimics a condensation reaction intermediate between malonyl-AcpP and a FabF bound hexanoic acid. However, future structural studies using stable malonyl-ACP or malonyl-CoA^61^ mimetics would likely enhance our understanding of the decarboxylation reaction mechanism.

**Figure 5:**
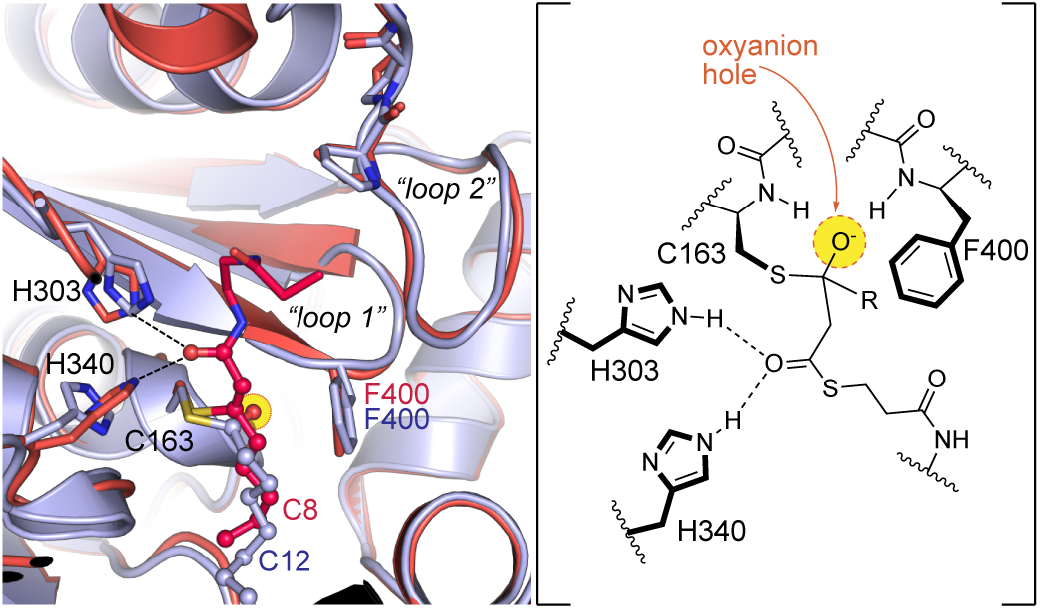
C8-AcpP=FabF mimics a condensation reaction intermediate. Superposition of C8Cl-AcpP-FabF and the C12-FabF (PDB: 2GFY) demonstrating that C8-AcpP=FabF mimics a condensation reaction intermediate between malonyl-AcpP and a FabF bound hexanoic acid. The carbonyl of the C8Cl crosslinker is coordinated to the two catalytic histidines (dashed lines). The overlaid C12-FabF structure shows that the oxyanion hole (yellow circle) would be occupied by the carbonyl of the bound acyl substrate. Enolate attack on the bound fatty acid would result in a tetrahedral intermediate stabilized by the oxyanion hole (shown as yellow circle). A 2D schematic of the putative condensation half-reaction tetrahedral intermediate is provided for comparison.

### C8Cl-AcpP=FabF delineates a unique Loop 2 interaction network

The position of loop 2 in the gate-closed conformation suggests that movement of loop 2 is required to allow loop 1 to access the gate-open conformation. The interaction network that stabilizes the loop 2 closed conformation (*apo-*FabF PDB: 2GFW) is mediated by a Ser271_(o)c-_His268_(H)N_ main-chain hydrogen bond, a Thr270-His268 side-chain interaction, and a Pro273-Tyr267 C-H π interaction (Figure 2A, Figure 6). Transition to the open conformation (C16-AcpP=FabF PDB: 6OKG) yields a distinct loop 2 interaction network, and interestingly, Thr270 swaps its interaction with His268 for Asp35 of the bound AcpP (Figure 2A, Figure 6).

**Figure 6:**
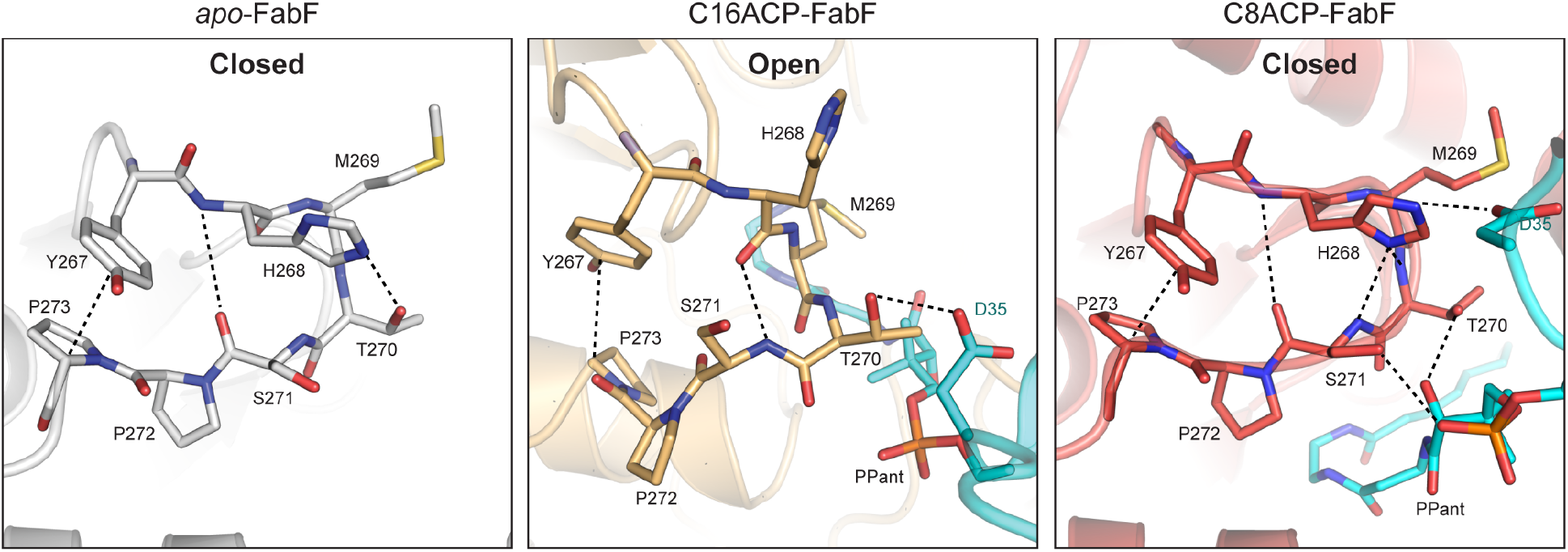
Loop 2 interaction networks. The *apo-*FabF, C16-AcpP-FabF, and C8-AcpP=FabF loop 2 interaction networks are distinct. Interactions between acyl-AcpP and loop 2 are present in the C8-AcpP=FabF (substrate bound, gate-closed) and C16-AcpP=FabF (substrate bound, gate-open) structures.

Analysis of the C8Cl-AcpP=FabF crosslinked complex reveals a loop 2 that, while in a similar conformation as seen in *apo-*FabF and C12-FabF, nonetheless uses a distinct interaction network. The Pro273-Tyr267 C–H π interaction is maintained, but His268 instead forms an interaction with Asp35 of AcpP and Thr270 and Ser271 form hydrogen-bonding interactions with the hydroxyl group of the PPant arm and the phosphoserine oxygen, respectively (Figure 6). Interestingly, similar loop 2 interaction networks are observed in the recently solved type II PKS ACP=KS-CLF complexes.^59^ Collectively, these findings suggest that unique AcpP—loop 2 interactions are present in both the closed and open conformations, providing additional evidence for an allosteric role in substrate binding and cargo delivery during KS gating events.

### The gating mechanism is important for the FabF condensation reaction

To confirm that FabF gating events are important for catalysis, we subjected the panel of FabF gating mutants used in our previous study^24^ to an HPLC-based kinetic assay^28^. These mutant kinetic assays aim to test gate function by blocking access to the gate open conformation (pocket-block), destabilizing the gate open conformation (destabilization), or limiting the flexibility of loop 1 (flex-reduction) (Figures S8-S11). Briefly, the assay monitors the rate of *holo-AcpP* formation in a reaction utilizing acyl-AcpP substrates as acyl-donors in the transacylation half-reaction while taking advantage of FabF’s relaxed specificity for malonyl-CoA (extender unit) for the condensation half-reaction (Figures S12, S13). We monitored the overall FabF condensation reaction for all previously tested gating mutants as well as an additional destabilization mutant, N404A, using both C6-AcpP and C12-AcpP substrates.

Using C6- or C12-AcpP, the apparent turnover rates of wt FabF in the tested conditions are 1.57 min^-1^ and 1.41 min^-1^, respectively, which are comparable to previously reported values^28^ (Table 1). Similar to a previous study by Zhang *et al*., the F400A impaired gate-removal mutant is 1-2% as active as wt FabF with both tested substrates.^32^ We then evaluated the activity for the two pocket-block mutants, G310M and G310F, where mutation of glycine to a bulky hydrophobic residue fills the pocket occupied by Phe400 when in the open conformation (Figure S9). These mutants demonstrated reduced condensation rates for both C6 and C12-AcpP substrates with an overall activity roughly 50-fold lower than that of wt FabF.

**Table 1:**
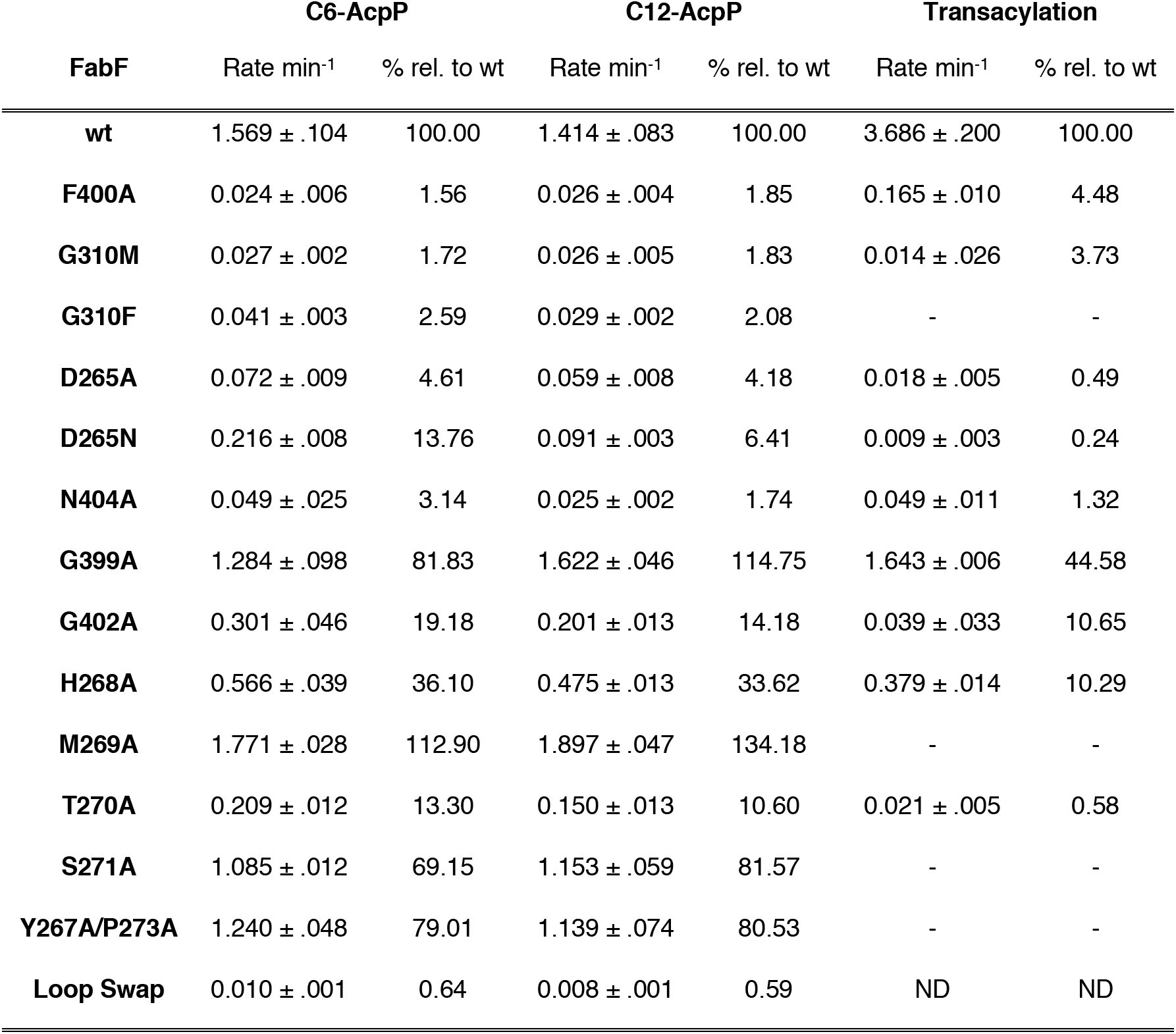
Quantification of condensation and transacylation rates of FabF mutants

The destabilization class of mutants was designed to disrupt the hydrogen-bonding network coordinated by Asp265, a highly conserved loop 2 residue that interacts with the backbone amides of loop 1 (in the gate-open conformation) and the side chain of Asn404 (Figure 2A, Figure S10.). The D265A mutant was 4.6% (0.07 min^-1^) and 4.8% (0.06 min^-1^) as active as wild-type with C6-AcpP and C12-AcpP, respectively. Interestingly, the more conservative D265N mutation was 13.8% (0.22 min^-1^) as active as wild-type with C6-AcpP but only 6.4% (0.09 min^-1^) as active as wild-type with C12-AcpP. To further validate the importance of this interaction network, we mutated the conserved Asn404 residue (Figure 2) to alanine and evaluated its activity with both substrates. As seen with D265A and D265N, the N404A mutant was a poor catalyst with rates 3.1% (0.05 min^-1^) and 1.7% (0.03 min^-1^) that of wt FabF with C6-AcpP and C12-AcpP, respectively.

The two flex-reduction mutants, G399A and G402A, were designed to inhibit gate function by limiting the conformational space available to the loop 1 GFGG motif (Figure S11). The G399A mutant was as fast as wt FabF, while rates for the G402A mutant were 19.2% (.301 min^-1^) and 11.0%(0.201 min^-1^) as active with C6- and C12-AcpP, respectively. These results are in line with our previous study as only G402A showed a reduction in crosslinking activity while G399A was as active as wt FabF. Taken together, our results indicate that all FabF gating mutants, with the exception of G399A, significantly decrease FabF catalyzed condensation rates.

### Transacylation rates of FabF gating mutants

Results from our assays demonstrate that FabF gating mutants affect the overall condensation reaction. We next attempted to verify that the FabF gating mechanism is important for the transacylation half-reaction. To address this question, we modified our HPLC assay to monitor the transfer of the C12 acyl chain from lauroyl-CoA to *holo-*ACP, effectively decoupling the transacylation and condensation reactions. Similar KS transferase assays have been used previously to assess the role of FabF active site residues on the transacylation step^32^ and to evaluate substrate specificity and cooperativity in the murine FAS^36^. Using these assay conditions, we were able to reliably determine transacylation rates for wt FabF and FabF gating mutants by monitoring the formation of C12-AcpP (Table 1, Figure S14).

Type I and II FAS KSs both possess the conserved Phe400 gating residue and only accept acyl-ACP substrates without β-carbon modifications (i.e. β-hydroxy-acyl-ACPs). The Phe400 gating residue was recently proposed to serve as a β-carbon sensor in the murine type I FAS KS domain.^36^ This model suggests that Phe400 is responsible for ensuring partially processed FAS intermediates are not accepted. However, previous work on type II FAS KS domains suggests that Phe400 minimally affects transacylation and primarily regulates the condensation step.^32^ Despite these previous reports, we determined a transacylation rate 4.5% that of wt for the F400A gate-deletion mutant (Table 1). Therefore, our results demonstrate that Phe400 is important for the transacylation half-reaction in addition to its well-established function in the condensation step.

The transacylation rate of the pocket block mutant, G310M, showed a similar reduction in transacylation activity as F400A, demonstrating that either blocking gate function or deleting the central gating residue decreases acyl-transfer rates. Interestingly, the destabilization mutants, D265A and D265N, distal to the KS active site had the largest decrease in activity with rates 200- and 300-fold lower than wt FabF, respectively (Table 1). Flex reduction mutants, G399A and G402A, showed a 2-fold and 10-fold drop in transacylation activity, respectively, suggesting that G399A does have a minor role during the transacylation step despite not reducing the overall condensation reaction rate.

Loops 1 and 2 were previously proposed to move in a coordinated manner to facilitate substrate processing during the transacylation half-reaction.^24^ Results from our condensation reaction assays demonstrate that blocking gate function significantly reduces overall KS-mediated substrate turnover. However, an argument could be made that these mutations alter malonyl-CoA binding or positioning of the catalytic histidines, thereby only disrupting the condensation half-reaction. Additionally, loop 1 coordinates the oxyanion hole for both half-reactions, suggesting that appropriate gate closure could regulate the condensation step as well. By implementing an assay that eliminates the condensation half-reaction, we demonstrated that the observed reductions in the overall KS condensation reaction for FabF gating mutants are correlated to significant reductions in transacylation rates, and therefore, the transacylation half-reaction.

### Importance of loop 2 residues for gate function

To quantitatively measure the importance of loop 2 for gate function, we subjected it to alanine scanning mutagenesis and tested all resulting soluble protein constructs using both our condensation and transacylation assays (Figure S8). In addition, we tested a FabF/FabB loop swap, where the loop 2 of FabB was used in place of FabF’s loop 2. The Y267A, P272A, and P273A mutants were insoluble, but we determined rates for alanine mutants of all remaining loop 2 residues as well as a Y267A/P273A double mutant. The overall KS condensation assay results indicate that only the conserved His268 and Thr270 (Figure 2B) are important for FabF activity. The H268A variant has a 3-fold reduction in activity, while the T270A mutation results in a 10-fold reduction. All other constructs are as active as wt FabF with the exception of the FabF/FabB loop swap variant, which has greater than a 150-fold drop in activity for both substrates (Table 1).

To determine if these changes in activity were related to the transacylation half-reaction, we determined transacylation rates for H268A, T270A, and the FabF/FabB loop swap variants. The H268A variant had a transacylation rate 10.3% that of wt FabF while the T270A variant had a rate 0.6 % that of wild-type. Interestingly, the transacylation rate for the FabF/FabB loop swap variant was too slow to be reliably measured (Table 1).

Results from the KS condensation and transacylation assays demonstrate that the conserved His268 and Thr270 residues, both of which interact with the bound acyl-AcpP in the closed and open states, respectively (Figure 6), are important for FabF activity. Mutation of these residues to alanine significantly decreases KS transacylation rates, indicating that they play a role in gate function and that acyl-AcpP binding might participate in triggering or stabilizing gate transitions via loop 2 interactions, as suggested by previous molecular dynamics (MD) simulations^24^. Interestingly, replacing FabF’s loop 2 with that of FabB results in an enzyme variant that is nearly inactive, providing additional evidence that loops 1 and 2 function in a coordinated manner, and that they may not be readily transferrable between different KS families. Therefore, additional elements, which may include ACP-KS PPIs, likely coevolve with loop 2 to fine-tune gate function. A dual loop gating mechanism suggests a dynamic and complex catalytic process, which is potentially coupled to ACP binding, chain-flipping of acyl cargo into the KS active site, and KS-substrate recognition. Therefore, future studies analyzing the putative gating loops in FabB, type I FAS, and PKS KS domains will be essential to conclusively address these latter hypotheses.

### Evaluating Type II FAS-Type I PKS loop 1 chimeras

Unlike FAS KSs, Type I PKS domains readily accept β-modified acyl-ACP substrates as well as a variety of α-branched extender units.^17–62^ Comparison of type I PKS KS domains and type II FAS KSs indicate that type I PKS KSs possess less bulky GISG and GVSG loop 1 motifs (Figure 2B) that could provide additional space to accommodate β-carbon modifications or α-branched extender units. Therefore, as in type II FAS, the putative gating machinery in type I PKS KS domains may be tuned to exert substrate specificity in either the transacylation or condensation half-reactions.

To determine how type I PKS gating elements would alter FabF activity and substrate specificity, we generated chimeric FabFs containing loop 1 elements from type I PKS, where the Phe400 gating residue was replaced with either isoleucine or valine. Additionally, we prepared two complete FabF-type I PKS KS loop 1 chimeras by mutating FabF’s GFGG motif to either GVSG (FabF-GVSG) or GISG (FabF-GISG). We first evaluated the overall condensation rates of our panel of loop 1 chimeras in assays containing either 0.25 mM or 1.0 mM malonyl-CoA (Table 2). The condensation rates for all tested mutants were significantly reduced as compared to wt FabF for both concentrations of malonyl-CoA. The F400V and F400I mutations turned over substrate with rates 4.4% and 5.5% that of wt FabF, respectively, at 0.25 mM malonyl-CoA and 4.6% and 6.9% that of wt FabF, respectively, at 1.0 mM malonyl-CoA. These mutants were roughly 3-fold and 10-fold more active than the gate deletion mutant, F400A, at 0.25 mM or 1.0 mM malonyl-CoA, respectively, indicating that reintroducing a gating residue recovers some activity. Condensation rates for the FabF-GVSG and FabF-GISG constructs were more significantly reduced with activities 1-2% that of wt FabF for both tested concentrations of malonyl-CoA.

**Table 2.** Quantification of condensation and transacylation rates of FabF gating mutants

| **FabF** | **.25 mM mCoA** |  | **1 mM mCoA** |  | **1 mM mmCoA** |  | **Transacylation** |  |
| --- | --- | --- | --- | --- | --- | --- | --- | --- |
|  | Rate min<sup>-1</sup> | % rel. wt | Rate min<sup>-1</sup> | % rel. wt | Rate min<sup>-1</sup> | % rel. wt | Rate min<sup>-1</sup> | % rel. wt |
| **wt** | 1.569 ± .104 | 100.00 | 5.011 ± .142 | 100.00 | 0.281 ± .056 | 100.00 | 3.686 ± .200 | 100.00 |
| **F400A** | 0.024 ± .006 | 1.56 | 0.027 ± .002 | 0.54 | 0.020 ± .009 | 7.14 | 0.165 ± .010 | 4.48 |
| **F400I** | 0.069 ± .003 | 4.44 | 0.232 ± .033 | 4.63 | 0.040 ± .002 | 14.33 | 0.371 ± .018 | 10.06 |
| **F400V** | 0.087 ± .013 | 5.53 | 0.346 ± .103 | 6.90 | 0.070 ± .034 | 25.03 | 0.400 ± .084 | 10.87 |
| **GISG** | 0.037 ± .009 | 2.34 | 0.084 ± .018 | 1.67 | 0.037 ± .007 | 13.09 | 0.016 ± .001 | 0.44 |
| **GVSG** | 0.023 ± .004 | 1.50 | 0.045 ± .005 | 0.90 | ND | ND | 0.009 ± .002 | 0.25 |

To ascertain if the measured reductions in activity were related to extender unity identity, we tested our FabF loop 1 chimeras with assays containing 1 mM methylmalonyl-CoA. We measured an apparent turnover number of 0.28 min^-1^ for wt FabF, an 18-fold reduction compared to assays with 1 mM malonyl-CoA (Table 2). Extending our analysis to the chimeric FabF constructs, we again found the condensation rates for all variants were decreased compared to wt FabF. Interestingly, the drop in activity for these mutants was less pronounced than in assays using malonyl-CoA. The F400V and F400I variants were 14% and 25% as active as wt FabF, respectively, and the GISG mutant was 13% as active. We were unable to reliably detect activity for the GVSG mutant in the tested conditions.

The overall reaction rates of FabF gating chimeras are slower than wt FabF with both malonyl-CoA and methylmalonyl-CoA, but it is unclear which half-reaction is more impacted by these mutations. Therefore, we evaluated the transacylation rates of our panel of FabF loop 1 chimeras (Table 2). The F400V and F400I variants have transacylation rates 10% that of wt FabF, again demonstrating that Phe400 is important for the transacylation half-reaction. Although, it is worth noting that the F400V and F400I constructs have transacylation rates 2.5-fold higher than F400A. The transacylation rates of the FabF-GVSG and FabF-GISG variants were found to be more significantly reduced, with activities 0.3% and 0.4% that of wt FabF, respectively. The more than 200-fold drop in transacylation activity for FabF-GVSG and FabF-GISG chimeras likely explains their low activities with either malonyl- or methylmalonyl-CoA.

In summary, replacement of Phe400 with either valine or isoleucine results in an enzyme more active than the F400A gate-removal mutant, but still 20-fold less active than wild-type with malonyl-CoA substrates. The transacylation rates for the F400V and F400I mutants are 10-fold lower than wt FabF, suggesting that both the transacylation and condensation half-reactions are affected by replacing Phe400 for type I PKS KS gating residues. Interestingly, in assays using methylmalonyl-CoA, the reduction in relative-to-wild-type activity for F400V, F400I, and FabF-GISG was less than for malonyl-CoA substrates. This result is especially pronounced for FabF-GISG, which was 13% as active as wt FabF with methylmalonyl-CoA and only 1.7% as active as wild-type with malonyl-CoA. The fact that FabF-GISG is only 10-fold less active than wild-type with methylmalonyl-CoA is quite interesting when considering that this mutant has a 225-fold reduction in transacylation rate. These results may indicate that loop 1 elements from other KS families are not readily compatible with the gating machinery in FabF. Indeed, the FabF loop 2 is divergent from that of type I PKS KSs and may not be tuned to coordinate gating transitions with GVSG and GISG loop 1 motifs (Figure 2B). This analysis is in agreement with our results from the FabF/FabB loop swap variant. Alternatively, although not mutually exclusive, the low transacylation rates of FabF loop 1 chimeras, especially for the FabF-GVSG and FabF-GISG constructs, could indicate that these loops are better suited for accepting acyl groups with β-substituents. Future KS engineering efforts should therefore consider the effect of loop 1 gating elements on both half reactions as the oxyanion hole coordinated by this loop is require to stabilize transition states during both half-reactions. Additionally, any mutations to loop 1 may need to be evaluated in relationship to loop 2 and substrate interactions in the acyl-binding pocket. Additional studies investigating the putative gating elements in related KS domains will be essential to addressing these unanswered questions.

## Conclusions

To better understand the KS reaction mechanism, we sought to further characterize our recently proposed KS gating mechanism with additional structural and catalytic investigations. The data reported herein are congruent with a model wherein acyl-ACP binding and substrate transfer is mediated by the correlated movement of two KS active site loops, loops 1 and 2. We provide biochemical evidence that proper gate function is required for KS-mediated substrate turnover, and we further demonstrate its importance for the transacylation half-reaction. Importantly, our structural and biochemical data indicate a potential role for loop 2 in regulating loop 1 gating transitions. We observe distinct interaction networks for FabF’s loop 2 in both the open and closed state, which include contacts to the bound acyl-AcpP molecule, suggesting an allosteric role connecting acyl-AcpP binding and substrate delivery to gate function. Loop 2 is conserved within, but not between KS families, and our kinetic data indicates that neither loop 2 nor loop 1 is readily transferrable between evolutionarily divergent KS families. Therefore, loop 2 may coevolve with loop 1, or other KS elements such as ACP-mediated PPIs, to fine-tune KS gate function.

Despite the central role elongating KSs play in FAS and PKS dependent metabolic pathways, there are only a few successful KS engineering attempts reported in the literature^37–41–58–63–64^. This challenge likely correlates with deficiencies in our fundamental understanding of the KS reaction mechanism. The manner in which KSs determine substrate preferences, for either the transacylation or condensation steps, remains poorly understood. Using a combination of chemical biology, structural biology, and biochemistry, we provide compelling evidence in support of the KS gating mechanism and an additional structural framework governing its role in catalysis. Given the conserved nature of these gating elements in both FAS and PKS, these series of studies reported herein provide part of the necessary catalytic foundation to design and carry out future KS mechanistic studies. Moreover, this work also accelerates the development of strategies to engineer metabolic pathways in a predictable manner.

## Acknowledgments

This work was supported by NIH GM095970. J.T.M. was supported by T32 GM832626. Portions of the work were also funded by the Arthur and Julie Woodrow Chair at the Salk Institute (to J.P.N.) and the Howard Hughes Medical Institute (to J.P.N.). The authors thank Dr. Ashay Patel for the concept and design of Figure 2A and Dr. Laetitia Misson for assistance editing the manuscript.

