## Supplementary Information for "Structure and mechanistic analyses of the gating mechanism of elongating ketosynthases"

**Table of contents:**


- **A. Biological protocols**
- **A1.** Protein expression and purification
- **A2.** Site directed mutagenesis
- **A3.** Loading AcpP with probe molecules (one-pot)
- **A4.** Preparation of AcpP=FabF crosslinked complexes
- **A5.** Crystallization, data collection, and refinement
- **A6.** Crosslinking assays with C16:1-*crypto*-ACP and FabF
- **A7.** Preparation of *holo*-AcpP and *acyl*-AcpP substrates
- **A8.** Ketosynthase condensation assay
- **A9.** Ketosynthase transacylation assay
- **A10.** Analysis of kinetic data.

- **B. Synthetic protocols**
- **B1.** Synthesis of chlorovinylacrylamide-pantetheinamide crosslinker
- **B2.** Synthesis of C16:1-α-bromo-pantetheinamide crosslinker
- **B3.** General synthetic methods
- **B4.** Synthesis of C16:1 α-bromo-pantetheinamide

- **C. Supplemental Figures and Tables**
- **Figure S1.** Synthetic route for the synthesis of the C16:1-α-bromo-pantetheinamide probe.
- **Figure S2.** One-pot loading of C16:1αBromo pantetheineamide probe and crosslinking gels.
- **Figure S3.** Time course crosslinking gels of C16:1-crypto-AcpP with FabF.
- **Figure S4.** C16:1-AcpP-FabF substrate omit maps.
- **Figure S5.** Overlay of apo-FabF (PDB: 2GFW), C16-AcpP=FabF (PDB: 6OKG), and C16:1-AcpP=FabF.
- **Figure S6.** Stereochemistry at crosslinker α-carbon.
- **Figure S7.** C8-AcpP-FabF substrate omit maps.
- **Figure S8.** SDS-PAGE gels of all FabF constructs used in kinetic assays.
- **Figure S9.** *Ra*tionale for FabF pocket block mutants.
- **Figure S10.** Rationale for FabF destabilization mutants.
- **Figure S11.** Rationale for FabF flex reduction mutants.
- **Figure S12.** Overview of the KS catalyzed condensation reaction assay
- **Figure S13.** C12-AcpP calibration curve used for quantifying concentrations in kinetic assays
- **Figure S14.** Overview of the KS catalyzed transacylation assay.
- **Table S1.** Table 1 X-ray crystallography data collection and refinement statistics
- **Table S2** Primers for site directed mutagenesis (SDM)

- **D. References**

### A. Biological protocols

#### A1. Protein expression and purification:

*E. coli* AcpP and *E. coli* FabF were grown according to previously reported protocols.[1,2]

#### A2. Site Directed Mutagensis

All site directed mutagenesis constructs were produced using the method developed Liu and Naismith.[3] The primers used to create mutations specifically for this study are provided in Table S2. Primers for all other FabF mutants not found in Table S2 were reported previously.[1] The FabF/FabB loop swap construct was created using the around-the-horn method from NEB with overhangs that coded for the loop 2 region to be inserted into FabF.

#### A3. Loading AcpP with probe molecules (one-pot method)

Preparation of all *crypto*-ACPs via the one-pot method was done as previously described.[1,2]

#### A4. Preparation of AcpP=FabF crosslinked complexes

Preparation of C16:1-crypto-AcpP=FabF and C8Cl-crypto-AcpP=FabF complexes was done as previously described.[1]

#### A5. Crystallization, data collection, and refinement

Crystals of the C16:1-crypto-AcpP=FabF and C8Cl-crypto-AcpP=FabF crosslinked complexes were grown by cross-seeding using crystals of C16-crypto-AcpP=FabF. 1.5 µL of crosslinked complex (8 mg•mL$^{-1}$) was mixed with 1.0 µL of the corresponding mother liquor and .5 µL of seed stock and the mixture was placed inverted over 500 µL of the well solution (hanging-drop method) at 6 °C. The same crystal conditions were used as previously described (26–30% PEG 8 K, 0.1 M sodium cacodylate pH 6.5, and 0.3 M NaOAc). All data were collected at the Advanced Light Source (ALS) synchrotron at Berkeley. Data were indexed using iMosflm[4] then scaled and merged using the aimless program from the CCP4 software suite[5]. Scaled and merged reflection output data was used for molecular replacement and model building in PHENIX[6]. Phases were solved using molecular replacement using C16-crypto-AcpP=FabF (PDB ID: 6OKG) as a search model with program Phaser from the PHENIX software suite. Model building was performed using Coot.[5] The parameter files for each of the covalently bonded 4'-phosphopantetheine analogs were generated using eLBOW in the PHENIX software suite[7]. Manually programmed parameter restraints using values provided by Jligand[8] were used to create the associated covalent bonds between 4'-phosphopantetheine to Ser36 and Cys92 during refinement.

#### A6. Crosslinking assays with C16:1-*crypto*-ACP and FabF

Each reaction was set up to contain 40 µM of crypto-AcpP, 20 µM of FabF, and buffer (25 mM Tris, 150 mM NaCl, 10% glycerol, 0.5 mM TCEP, pH depends), as previously described.[1] All sample stocks were originally in pH 7.4 buffer. Prior to the pH 6.0 and pH 7.0 crosslinking assays, samples were buffer exchanged via mini spin column to the corresponding pH. To stop the reaction at the designated time point, 2.5 µL of the reaction solution was mixed with 10 µL of 3X SDS dye. Samples were analyzed by 12% SDS-PAGE.

#### A7. Preparation of holo-AcpP and acyl-AcpP substrates

Preparation of holo-ACP and acyl-AcpP substrates was performed as previously described.[1,2]

#### A8. Ketosynthase condensation assay

Ketosynthase condensation assays were carried out as previously described.[2] In assays run with 1000 µM malonyl-CoA, concentrations of all other assay components were held constant. Scouting reactions to evaluate effective time points for accurate rate rate determination were performed for each FabF mutant before running assays in triplicate. All assays were performed in biological triplicate and the reported error is the standard deviation on the mean.

**A9. Ketosynthase transacylation assay**

The FabF transacylation assay mixture contained 50 mM sodium phosphate (pH 6.8), 50 mM NaCl, 0.5 mM TCEP, 150 µM lauroyl-CoA, 25 µM *holo*-AcpP (wt), and 1 µM FabF (0 µM FabF for the control reaction) at 28 °C. Twenty microliters of each reaction mixture was quenched by the addition of 10 µL of 1.0% TFA (final TFA concentration of 0.33%) at five different time points to ensure accurate linear regression with at least four data points. Product formation was monitored by chromatographic separation using an Agilent 1100 series high-performance liquid chromatography (HPLC) instrument equipped with an Ascentis Express Peptide ES-18 column (15 cm × 4.6 mm, 2.7 µm) with a 1 mL/min flow rate and a detection wavelength of 210 nm. A gradient method was used [A: doubly distilled H2O with 0.1% (v/v) TFA; B: HPLC grade CH3CN with 0.1% (v/v) TFA]: 25% B, 0 min; 25% B, 2 min; 75% B, 12 min; 95% B, 13 min; 95% B, 15 min; 25% B, 17 min; 25% B, 20 min. Chromatographic traces of the wt-FabF transacylation assay can be found in Figure S13. C12-AcpP concentrations were calculated from a standard curve and used to determine initial reaction rates. All assays were performed in biological triplicate and the reported error is the standard deviation on the mean.

**A10. Analysis of kinetic data.** Kinetic rate data were analyzed with excel and Prism. Data points from were fit by linear regression to give rates for substrate turnover. The averages of triplicates were calculated, and the standard deviation is reported for each biological triplicate.

### B. Synthetic protocols

#### B1. Synthesis of C8Cl-chlorovinylacrylamide-pantetheinamide crosslinker

Preparation and synthesis of the C8Cl-chlorovinylacrylamide-pantetheinamide crosslinker was done as previously reported.[1,9]

#### B2. Synthesis of C16:1-α-bromo-pantetheinamide crosslinker

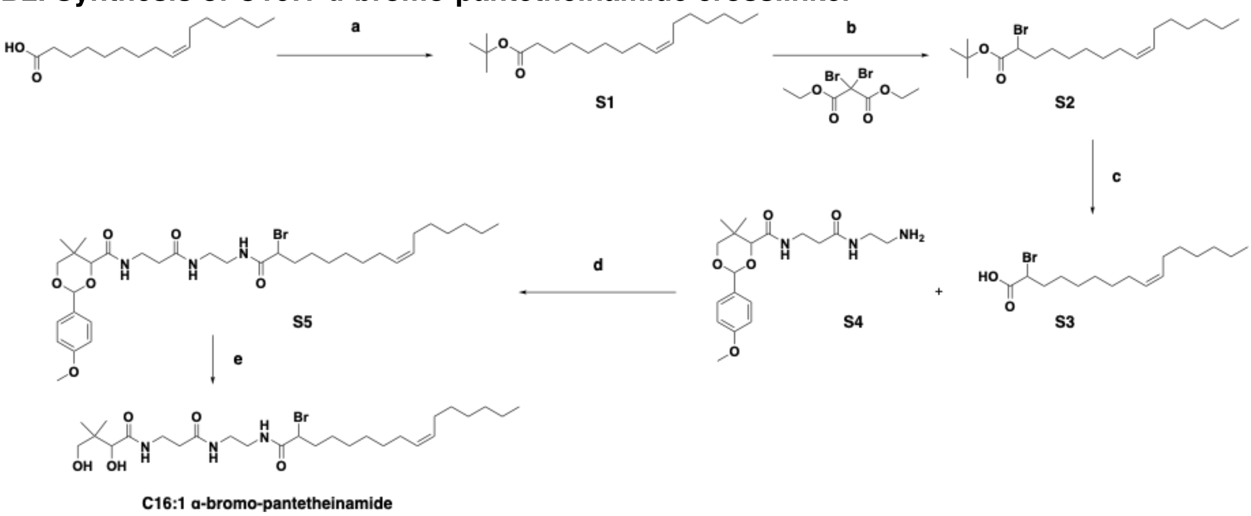


**Scheme S1. Synthetic route to C16:1 α-bromo-pantetheinamide.** *Reagents and conditions*: [a] $(BOC)_2O$, DMAP, DIEA, $CH_2Cl_2$, rt, 38%; [b] n-BuLi, DIPA, THF, -78 °C 63%; [c] 40% TFA/$CH_2Cl_2$, quantitative yield; [d] EDC, DMAP, $CH_2Cl_2$, rt, 40%; [e] 80% aq. acetic acid, rt, 37%.

**B3. General Synthetic Methods.** All commercial reagents were used as provided unless otherwise indicated. Compound **S4** is a known compound. This compound was prepared according to published literature procedures.[10] All reactions were carried out under an atmosphere of nitrogen in dry solvents with oven-dried glassware and constant magnetic stirring unless otherwise noted. $^1$H-NMR spectra were recorded at 500 MHz. $^{13}$C-NMR spectra were recorded at 125 MHz on JEOL NMR spectrometers and standardized to the NMR solvent signal as reported by Gottlieb.[11] Multiplicities are given as s = singlet, d = doublet, t = triplet, q = quartet, p = pentet, dd = doublet of doublets, ddd = doublet of doublet of doublets, dt = doublet of triplets, tt = triplet of triplets, dq = doublet of quartets, dp = doublet of pentets, qt = quartet of triplets, ddt = doublet of doublet of triplets, ddq = doublet of doublet of quartet, and m = multiplet using integration and coupling constant in Hertz. TLC analysis was performed using Silica Gel 60 F254 plates (Merck) and visualization was accomplished with ultraviolet light (λ = 254 nm) and/or the appropriate stain [phosphomolybdic acid, iodine, ninhydrin, and potassium permanganate]. Silica gel chromatography was carried out with SilicaFlash F60 230-400 mesh (Silicycle), according to the method of Still.[12]

#### B4. Synthesis of C16:1 α-bromo-pantetheinamide.

**Chemical Synthesis of S1.** Compound name in bold refers to the structures shown in Scheme S1.

**Compound S1**

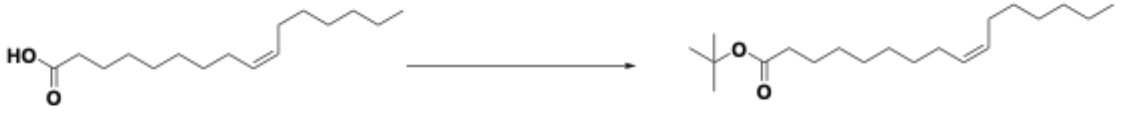

S1

$(BOC)_2O$ (197 mg, 0.9 mmol), DIEA (112 mg, 0.86 mmol), and DMAP (29 mg, 0.24 mmol) were added to a solution of the starting material, palmitoleic acid, (200 mg, 0.79 mmol) in $CH_2Cl_2$ (5 mL). The solution was stirred at room temperature overnight. The reaction mixture was diluted with $CH_2Cl_2$ and washed with $H_2O$ three times, dried over $Na_2SO_4$, filtered, and evaporated to dryness. The resulting red oil was purified by flash chromatography (hexane:EtOAc = 9:1) to afford compound **S1** as an oil (92 mg, 38%). $^1H$ NMR (500 MHz, $CDCl_3$): δ NMR

**Chemical synthesis of S2.**
**Compound S2**

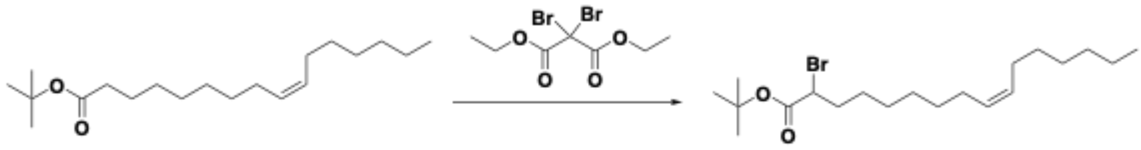

S1 S2

A 1.6 M solution of *n*-BuLi in hexane (0.6 mL, 0.96 mmol) was added to a solution of DIPA (0.15 mL, 1.04 mmol) in THF (2 mL) at -78 °C under inert conditions and stirred for 15 minutes to prepare LDA *in situ*. The compound **S1** (270 mg, 0.87 mmol) was dissolved in THF (3 mL) in a different vial at -78 °C and LDA (0.13 mL, 0.96 mmol) was cannulated into the solution. The reaction mixture was stirred at room temperature for 1 hour. In a different vial, diethyl dibromomalonate (0.6 mL, 2.17 mmol) was dissolved in THF (3 mL) and cooled to -78 °C and this solution was cannulated into the mixture containing the compound **S1** and LDA, which was then stirred for 2 h at -78 °C. The reaction mixture was quenched with 2% HCl, diluted with brine and hexane, washed with $H_2O$ three times, dried over $Na_2SO_4$, filtered, and evaporated to dryness. The resulting oil was purified by flash chromatography (hexane:DCM gradient from 10:1 to 8:1 to 6:1) to give compound **S2** as an oil (213 mg, 63%). $^1H$ NMR (500 MHz, $CDCl_3$): δ NMR

**Chemical synthesis of S5.**
**Compound S5**

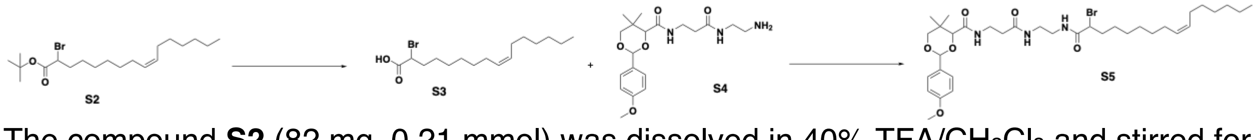

The compound **S2** (82 mg, 0.21 mmol) was dissolved in 40% TFA/$CH_2Cl_2$ and stirred for 1 h at room temperature. Solvent was evaporated by blowing air into the vial and reaction mixture was azeotrope with cyclohexane to give compound **S3** in quantitative yield. Compound **S4** (80 mg, 0.21 mmol), EDC (39 mg, 0.25 mmol), and DMAP (5 mg, 0.04 mmol) were added to a solution of Compound **S3** (70 mg, 0.21 mmol) in DMF (5 mL). The solution was stirred at room temperature overnight. The mixture was diluted with water and extracted with DCM three times. The combined organic layers were washed with $H_2O$ three times and then washed with brine, dried over $Na_2SO_4$, filtered, and evaporated to dryness. The resulting oil was purified by flash chromatography (DCM:methanol = 9:1, 1% TEA) to give compound **S5** as an oil (58 mg, 40%). $^1$H NMR (500 MHz, $CDCl_3$): δ 7.43 (d, *J* = 8.8 Hz, 2H), 7.06 (m, 1H), 7.01 (m, 1H), 6.92 (d, *J* = 8.8 Hz, 2H), 6.56 (bs, 1H), 5.47 (s, 1H), 5.34 (m, 2H), 4.23 (dt, *J* = 8.4, 5.2 Hz, 1H), 4.10 (m, 1H), 3.82 (s, 3H), 3.69 (q, *J* = 11.2 Hz, 2H), 3.56 (m, 2H), 3.39 (m, 4H), 2.44 (t, *J* = 6.2 Hz, 2H), 2.05 (m, 6H), 1.27 (m, 16H), 1.10 (m, 6H), 0.88 (t, *J* = 6.9 Hz, 3H). $^{13}$C NMR (125 MHz, $CDCl_3$) *δ* 172.0, 170.1, 169.9, 160.4, 130.2, 129.8, 127.7, 113.9, 101.5, 83.9, 78.6, 55.5, 51.4. 40.7, 39.5, 36.6, 35.8, 35.1, 33.3, 31.9, 31.7, 29.9, 29.2, 29.1, 28.9, 27.4, 27.4, 27.3, 22.8, 19.3, 14.3. HRMS (ESI) *m/z* $[M+Na]^+$ Calculated for $C_{35}H_{56}BrN_3O_6Na$: 716.3250; Found 716.3248.

**Chemical synthesis of C16:1 α-bromo-pantetheniamide**
**C16:1 α-bromo-pantetheinamide**

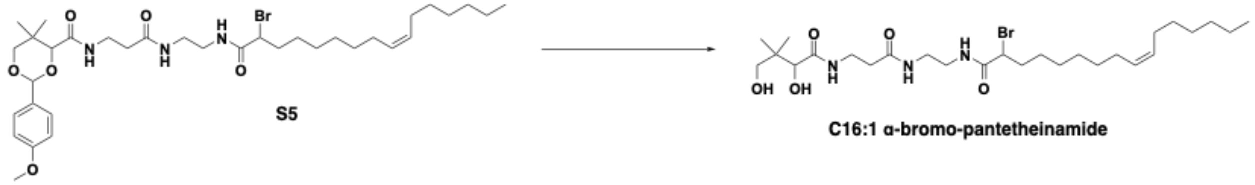


Compound **S5** (50 mg, 72 mmol) was dissolved in aqueous 80% acetic acid. The reaction mixture was stirred at room temperature overnight. The solution was dried by blowing air into the vial. The resulting oil was purified by flash chromatography (DCM:methanol = 5:1) to give the final product as an oil (15 mg, 37%). $^1$H NMR (500 MHz, $CDCl_3$): *δ* 7.00 (bs, 1H), 6.63 (bs, 1H), 6.48 (bs, 1H), 5.35 (m, 2H), 4.26 (m, 1H), 4.03 (d, *J* = 10.6 Hz, 1H), 3.70 (m, 1H), 3.50 (m, 5H), 3.36 (m, 2H), 2.45 (m, 2H), 2.34 (t, *J* = 7.6 Hz, 1H), 2.00 (m, 6H), 1.40 (t, *J* = 7.4 Hz, 1H), 1.27 (m, 16H), 1.02 (s, 3H), 0.93 (s, 3H), 0.88 (m, 3H). $^{13}$C NMR (125 MHz, $CDCl_3$) *δ* 174.1, 173.9, 170.6, 130.3, 129.7, 76.4, 71.2, 51.2, 40.4, 39.4, 36.3, 35.6, 32.3, 32.1, 31.9, 29.9, 29.2, 28.9, 27.4, 27.4, 27.3, 23.6, 22.8, 21.9, 20.7, 14.3. HRMS (ESI) *m/z* $[M+Na]^+$ Calculated for $C_{27}H_{50}BrN_3O_5Na$: 598.2832; Found 598.2826.

### C. Supplementary Figures and Tables

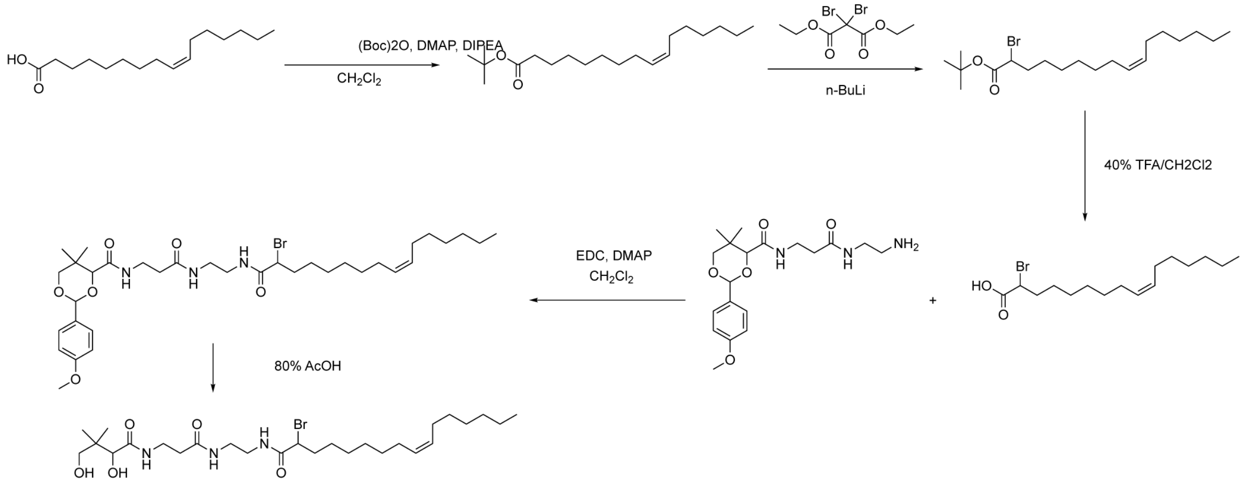


**Figure S1:** *Synthetic route for the synthesis of the C16:1-α-bromo-pantetheinamide probe.*

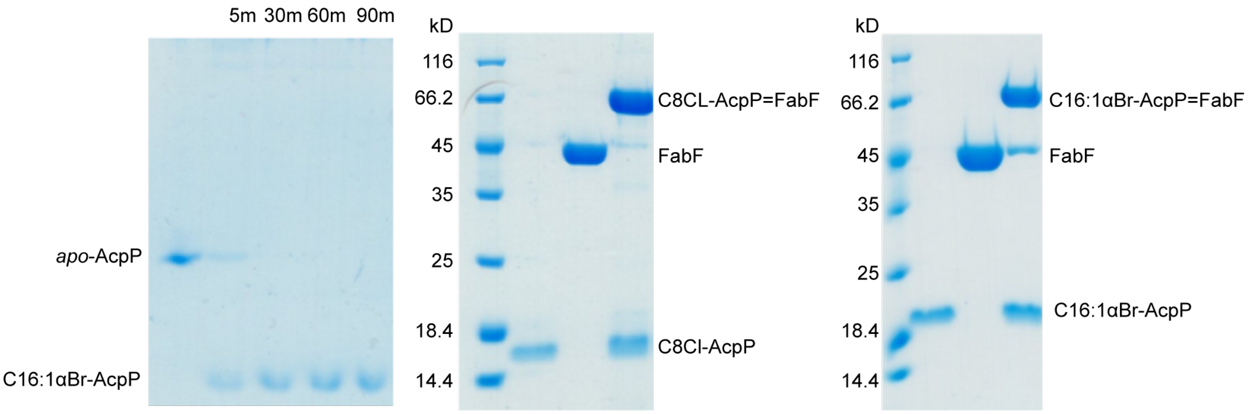


**Figure S2:** *One-pot loading of C16:1αBromo pantetheineamide probe and crosslinking gels.* A time course one-pot of *apo*-AcpP with the C16:1αBr-pantetheinamide probe (left). Scaled up crosslinking for sample preparation of C8Cl-AcpP=FabF for crystal trials (middle). Scaled up crosslinking for sample preparation of C16:1αBr-AcpP=FabF for crystal trials (right).

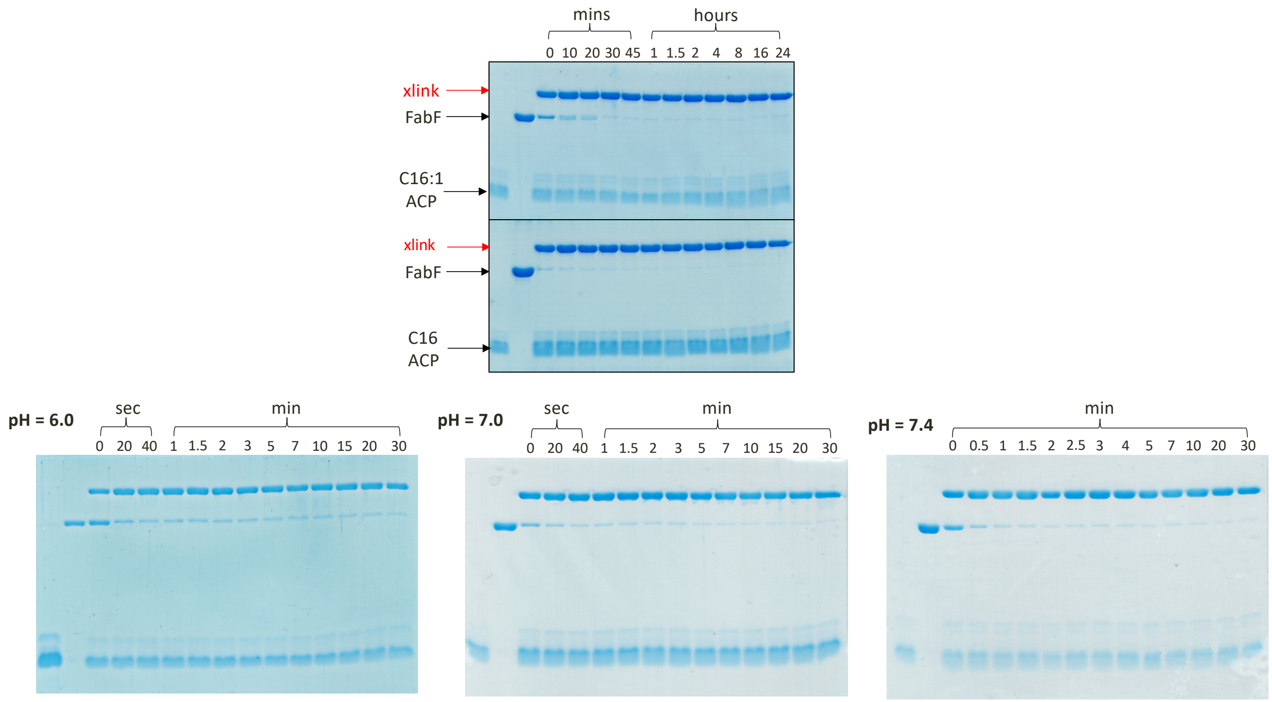


**Figure S3:** *Time course crosslinking gels of C16:1-crypto-AcpP with FabF.* The two gels on the top represent time-course crosslinking gels of C16:1- and C16-*crypto*-AcpP with FabF. The bottom three gels represent time-course crosslinking analysis of C16:1-*crypto*-AcpP with FabF at three different pHs (6.0, 7.0, and 7.4).

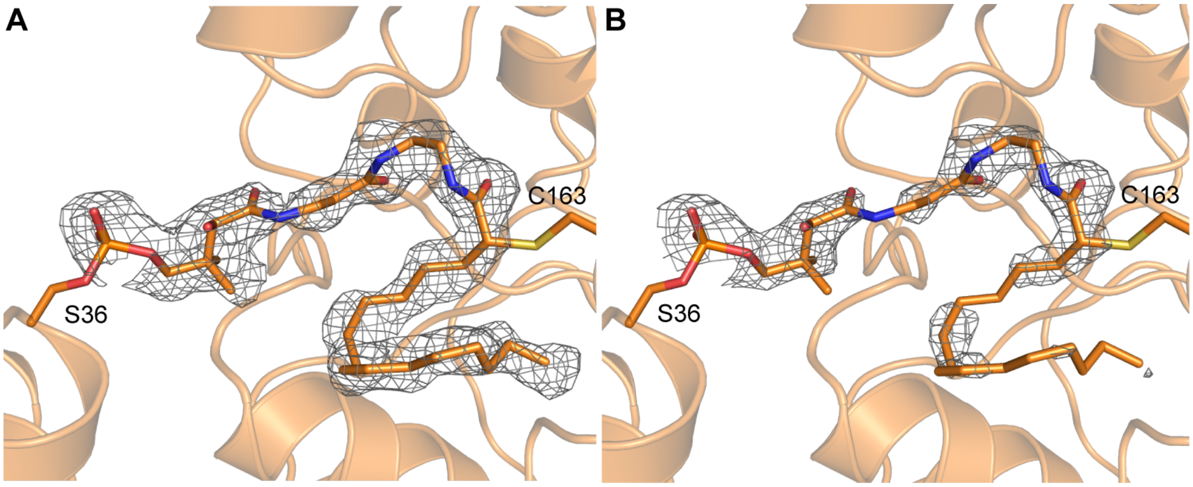


**Figure S4:** *C16:1-AcpP-FabF substrate omit maps.* **A**) Fo-Fc polder omit map calculated by omitting PPant crosslinker and bulk solvent from the region defined by a 5 Å solvent exclusion radius. The Fo-Fc polder map is contoured at 3.0 σ and rendered using a carve radius of 1.6 Å. **B)** Standard Fo-Fc omit map (no bulk solvent exclusion from omitted region) contoured at 3.0 σ and rendered using a carve radius of 1.6 Å.

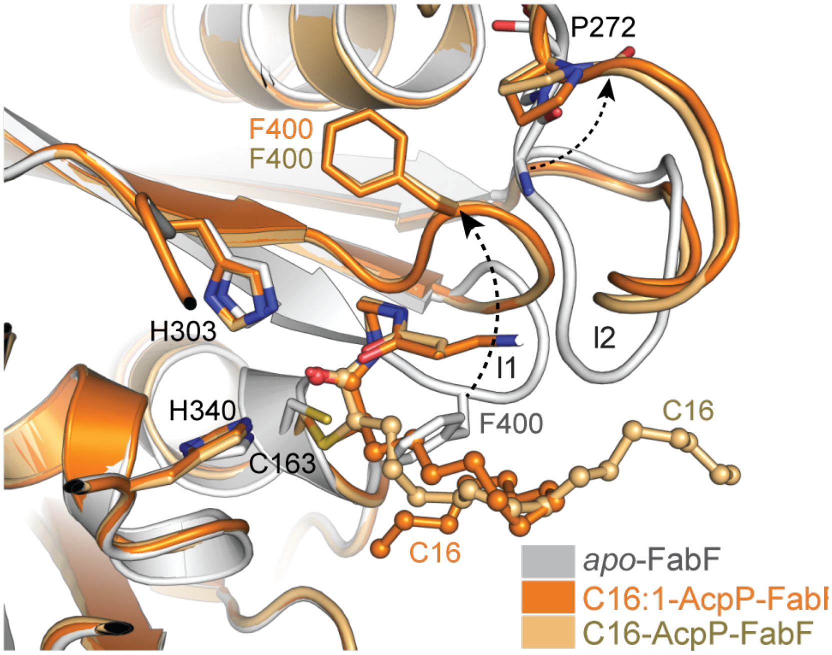


**Figure S5:** *Overlay of apo-FabF (PDB: 2GFW), C16-AcpP=FabF (PDB: 6OKG), and C16:1-AcpP=FabF.* The gating loops (I1 and I2) are found in the open conformation for both C16-AcpP=FabF and C16:1-AcpP=FabF. The arrows show the proposed trajectory of loop movement from the closed state (*apo*-FabF) to the open state. The fatty acid substrate mimetic of the C16 and C16:1 crosslinking probes are shown in ball and stick.

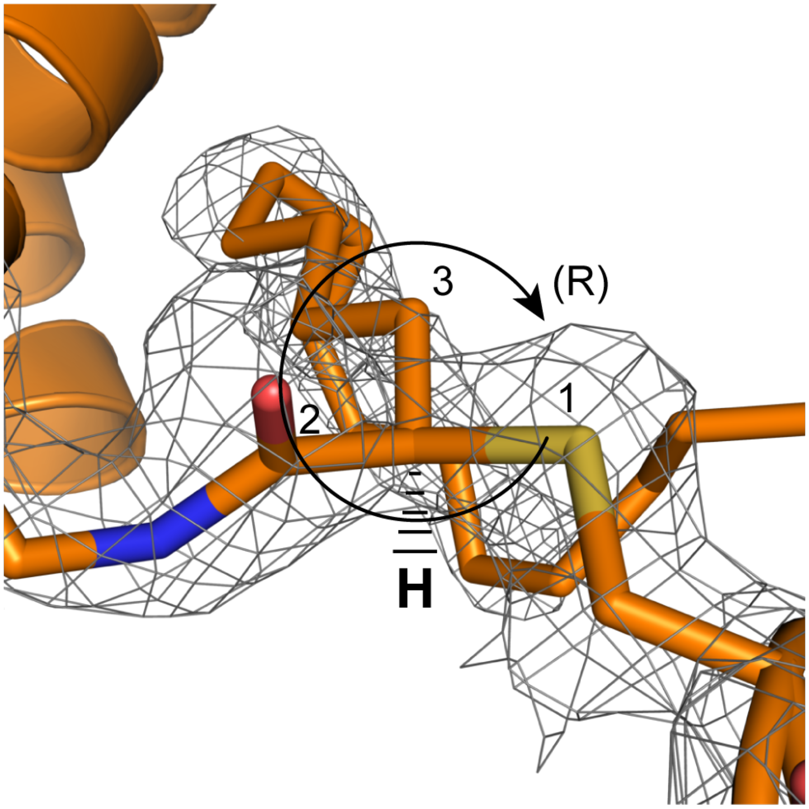


**Figure S6:** *Stereochemistry at crosslinker α-carbon.* The 2Fo-Fc electron density map show density for the thiovynyl crosslink site. The map is contoured at 1.0 σ and rendered using a carve radius of 1.6 Å. A hydrogen atom is added below to demonstrate that the substituent priorities (Cahn-Ingold-Prelog), based on our refinements, are arranged in the *R* configuration.

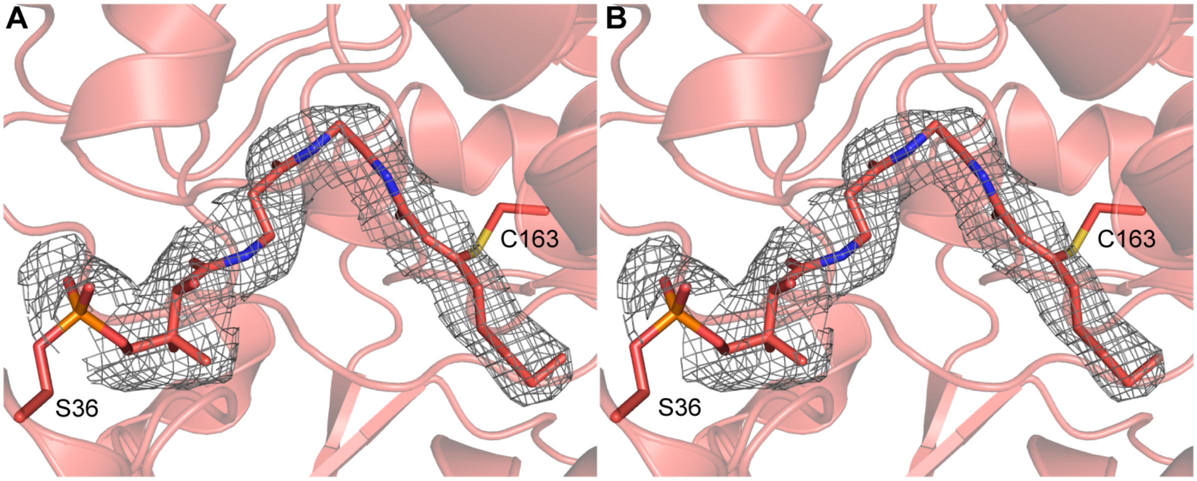


**Figure S7:** *C8-AcpP-FabF substrate omit maps.* **A**) Fo-Fc polder omit map calculated by omitting PPant crosslinker and bulk solvent from the region defined by a 5 Å solvent exclusion radius. The Fo-Fc polder map is contoured at 3.0 σ and rendered using a carve radius of 1.6 Å. **B)** Standard Fo-Fc omit map (no bulk solvent exclusion from omitted region) contoured at 3.0 σ and rendered using a carve radius of 1.6 Å.

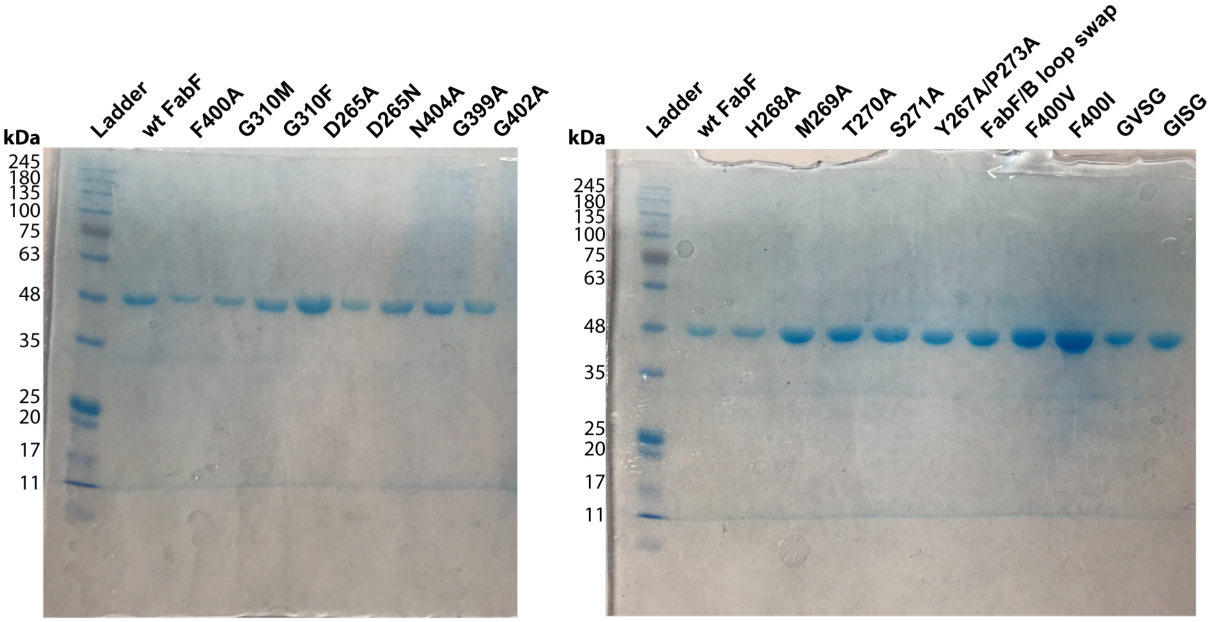


**Figure S8:** *SDS-PAGE gels of all FabF constructs used in kinetic assays*.

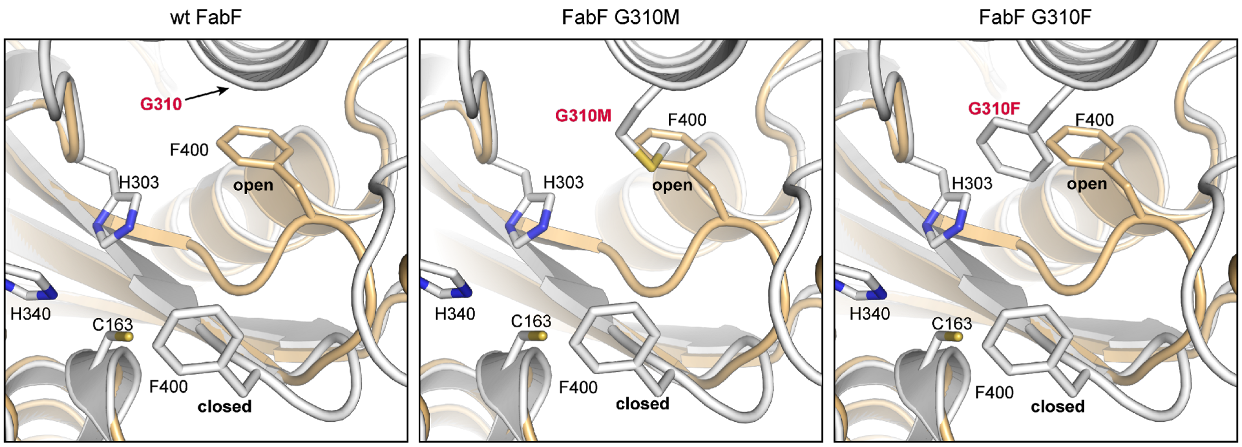


**Figure S9:** *Rationale for FabF pocket block mutants.* The open conformation of loop 1 positions Phe400 into a previously unoccupied pocket. Inserting bulky amino acids into this position should block access to the gate-open conformation, thereby limiting gate function.

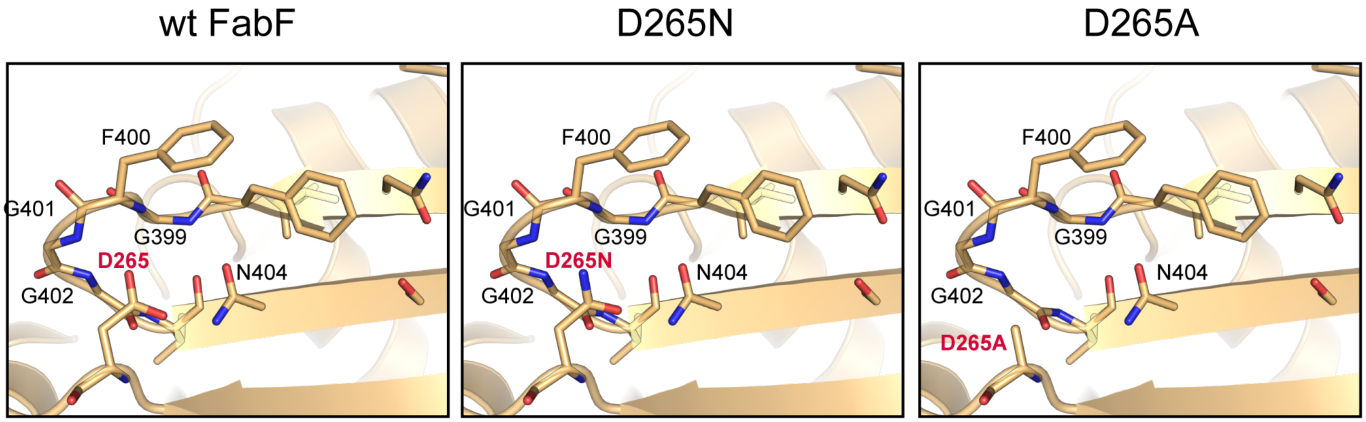


**Figure S10:** *Rationale for FabF destabilization mutants.* The open conformation of loop 1 is stabilized by a hydrogen bonding interaction network coordinated by the conserved D265 residue from loop 2. Mutation of this negatively charged aspartate residue for polar or non-charged amino acids should disrupt this network, thereby destabilizing the open conformation and inhibiting gate function.

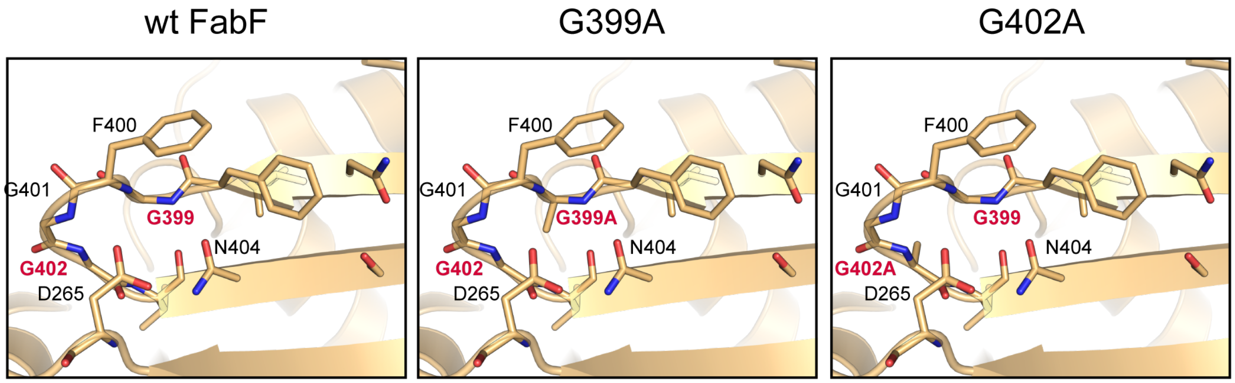

**Figure S11:** *Rationale for FabF flex reduction mutants.* Movement of loop 1 to the gate open conformation likely requires the conformational freedom of the conserved glycine residues from the GFGG motif. Mutation of these glycine residues (G399 and G402A) to alanine should limit the allowable backbone dihedral conformations, thus inhibiting gate function by hindering movement to the open conformation.

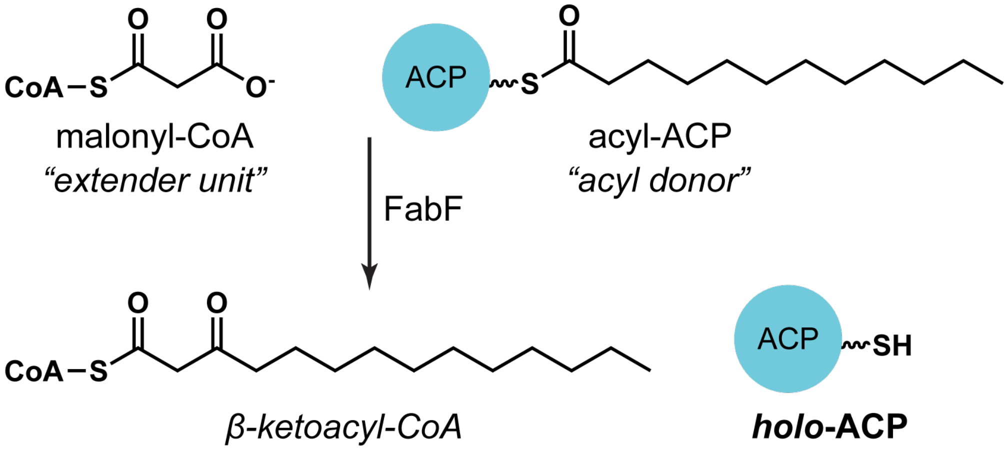


**Figure S12:** *Overview of the KS catalyzed condensation reaction assay*

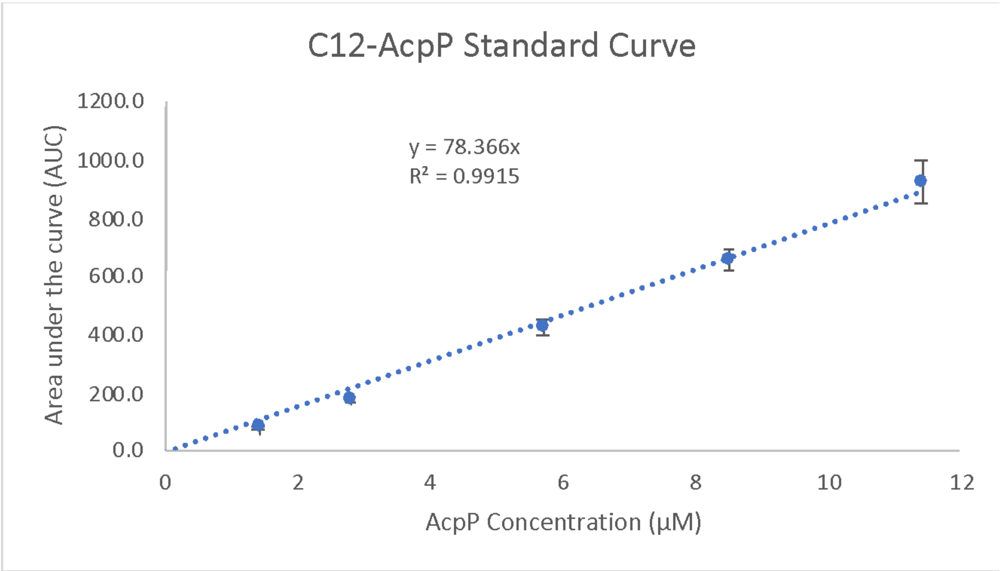


**Figure S13:** *C12-AcpP calibration curve used for quantifying concentrations in kinetic assays*

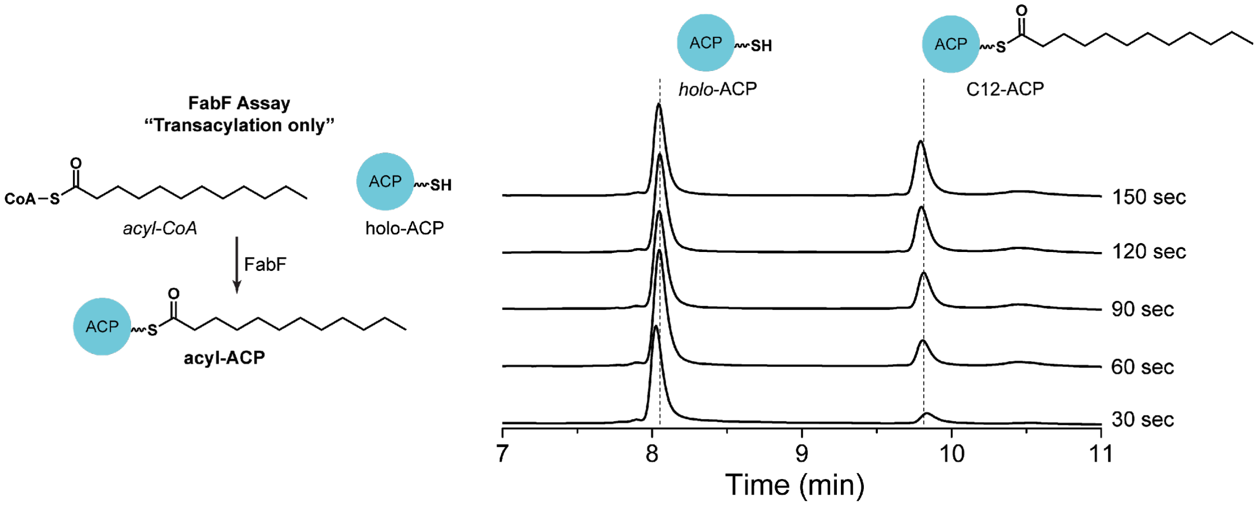


**Figure S14:** *Overview of the KS catalyzed transacylation assay.* The transacylation assay utilizes an acyl-CoA donor and holo-ACP acceptor and monitors the formation of the resulting acyl-ACP. A stacked HPLC chromatogram demonstrates the abilitiy to monitor acyl-ACP formation over time, thereby allowing accurate rates to be determined by this assay format.

**Table S1:** *Table 1 x-ray crystallography data collection and refinement statistics*

| | **C8-AcpP-FabF** | **C16:1-AcpP-FabF** |
|---|---|---|
| Wavelength | 1 | 1 |
| Resolution range | 48.4 - 2.65 (2.745 - 2.65) | 54.17 - 2.0 (2.071 - 2.0) |
| Space group | P 21 21 21 | P 43 21 2 |
| Unit cell | 83.782 92.136 113.746 90 90 90 | 86.9499 86.9499 114.48 90 90 90 |
| Total reflections | 364173 (37926) | 718278 (43525) |
| Unique reflections | 26219 (2554) | 29648 (2973) |
| Multiplicity | 13.9 (14.8) | 24.2 (16.4) |
| Completeness (%) | 99.92 (100.00) | 99.96 (99.93) |
| Mean I/sigma(I) | 7.49 (1.72) | 8.12 (0.69) |
| Wilson B-factor | 36.83 | 24.31 |
| R-merge | 0.3958 (1.907) | 29.02 (-152.1) |
| R-meas | 0.411 (1.974) | 32.1 (-172.2) |
| R-pim | 0.11 (0.5101) | 11.89 (-71.65) |
| CC1/2 | 0.989 (0.713) | 0.241 (0.228) |
| CC* | 0.997 (0.912) | 0.623 (0.61) |
| Reflections used in refinement | 26203 (2554) | 30316 (2972) |
| Reflections used for R-free | 1299 (137) | 1991 (196) |
| R-work | 0.2221 (0.3027) | 0.1671 (0.2756) |
| R-free | 0.2664 (0.3773) | 0.2068 (0.3442) |
| CC(work) | 0.946 (0.833) | 0.005 (0.039) |
| CC(free) | 0.901 (0.752) | 0.007 (-0.050) |
| Number of non-hydrogen atoms | 7266 | 3973 |
| macromolecules | 7070 | 3643 |
| ligands | 62 | 46 |
| solvent | 134 | 284 |
| Protein residues | 973 | 488 |
| RMS(bonds) | 0.004 | 0.004 |
| RMS(angles) | 0.97 | 1.02 |
| Average B-factor | 47.16 | 32.36 |
| macromolecules | 47.42 | 31.61 |
| ligands | 44.53 | 51.18 |
| solvent | 34.98 | 38.92 |
| Number of TLS groups | 24 | 16 |

**Table S2:** *Primers for site directed mutagenesis (SDM)*

| **Name** | **Sequence** |
|---|---|
| **FabF-F400V-F** | *TTCGGCGTTGGTGGCACTAATGGTTCTTTGATCTTTAAAAAGATC* |
| **FabF-F400V-R** | GCCACCAACGCCGAAGGAGTTACACAGAGTGTATTCCATTC |
| **FabF-F400I-F** | *TTCGGCATTGGTGGCACTAATGGTTCTTTGATCTTTAAAAAGATC* |
| **FabF-F400I-R** | GCCACCAATGCCGAAGGAGTTACACAGAGTGTATTCCATTC |
| **FabF-GVSG-F** | *TTCGGCGTTTCTGGCACTAATGGTTCTTTGATCTTTAAAAAGATC* |
| **FabF-GVSG-R** | GCCAGAAACGCCGAAGGAGTTACACAGAGTGTATTCCATTC |
| **FabF-GISG-F** | *TTCGGCATTTCTGGCACTAATGGTTCTTTGATCTTTAAAAAGATC* |
| **FabF-GISG-R** | GCCAGAAATGCCGAAGGAGTTACACAGAGTGTATTCCATTC |
| **FabF-N404A-F** | TTCGGTGGCACTGCGGGTTCTTTGATCTTTAAAAAGATCTAA |
| **FabF-N404A-R** | CGCAGTGCCACCGAAGCCGAAGGAGTTACACAGAGT |
| **FabF-Y267A-P273A-F** | GCTCATATGACGTCACCGGCAGAAAATGGCGCAGGCGCAGCTC |
| **FabF-Y267A-P273A-F** | TGCCGGTGACGTCATATGAGCAGCATCGCTGCTCATACCAAAGCC |
| **FabF-H268A-F** | TATGCGATGACGTCACCGCCAGAAAATGGCGCAGGCGCA |
| **FabF-H268A-R** | CGGTGACGTCATCGCATAAGCATCGCTGCTCATACCAAAGCC |
| **FabF-M269A-F** | TATCATGCGACGTCACCGCCAGAAAATGGCGCAGGCGCA |
| **FabF-M269A-R** | CGGTGACGTCGCATGATAAGCATCGCTGCTCATACCAAAGCC |
| **FabF-S271A-F** | TATCATATGACGGCGCCGCCAGAAAATGGCGCAGGCGCA |
| **FabF-S271A-R** | CGGCGCCGTCATATGATAAGCATCGCTGCTCATACCAAAGCC |
| **FabF-FabB-LoopSwap-F** | GTTGCTCCGTCTGGCGCAGGCGCAGCTCTG |
| **FabF-FabB-LoopSwap-R** | CATGTCTGCACCATCGCTGCTCATACCAAAGCCG |
